# Craniofacial studies in chicken embryos confirm the pathogenicity of Frizzled2 variants associated with Robinow syndrome

**DOI:** 10.1101/2023.11.07.565956

**Authors:** Shruti S. Tophkhane, Katherine Fu, Esther M. Verheyen, Joy M. Richman

**Affiliations:** Life Sciences Institute and Faculty of Dentistry, University of British Columbia, Vancouver, Canada; Department of Molecular Biology and Biochemistry, Centre for Cell Biology, Development and Disease, Simon Fraser University, Burnaby, Canada

**Keywords:** Wnt signaling, intramembranous ossification, chondrogenesis, micromass cultures, luciferase assays, RCAS virus

## Abstract

Robinow syndrome (RS) is a rare disease caused by mutations in seven WNT pathway genes. Features include craniofacial widening and jaw hypoplasia. We used the chicken embryo to test two autosomal dominant RS (ADRS) missense *FZD2* variants on the frontonasal mass, the affected region in RS. The wild-type (wt) and variant h*FZD2* inhibited beak ossification. The bone hypoplasia was possibly mediated by decreased levels of WNT and BMP pathway genes. In primary cultures, h*FZD2* variants inhibited chondrogenesis, increased nuclear shuttling of β-catenin and increased expression of TWIST1, both known to suppress chondrogenesis. In luciferase reporter assays, proteins coding for *1301G>T* and *425C>T* FZD2 variants weakly activated canonical WNT reporter and dominantly interfered with wt*FZD2*. In the JNK-PCP WNT pathway luciferase assay, only the *425C>T* showed a loss-of-function. The 1301G>T variant presumably acts through a JNK-independent pathway. This is the first study to demonstrate that the ADRS-*FZD2* missense variants cause craniofacial and WNT signaling defects. Frontonasal mass width is increased by both h*FZD2* variants which sheds light on the ontogeny of the broad facial features seen in individuals with RS.

**Summary Statement:** Gain-of-function studies on *FZD2* missense variants associated with Robinow syndrome led to increased facial width, altered Wnt signaling and inhibition of beak skeletogenesis in chicken embryos.

## INTRODUCTION

Multiple syndromes include craniofacial dysmorphology as part of the clinical presentation. Studying the syndrome causing mutations sheds light on gene function in many aspects of facial morphogenesis, patterning and differentiation. In this study, we focused on Robinow syndrome (RS), a rare skeletal dysplasia syndrome (1:500,000 live births) that primarily affects the skeleton of the face and limbs (Mazzeu and Brunner, 2020). RS was first reported in 1969 in a family with dwarfism and wide spaced eyes or hypertelorism (Robinow et al., 1969). Approximately 250 cases of RS have been reported so far in the literature (Schwartz et al., 2021; Suresh, 2008). Interestingly, the seven genes associated with the pathogenesis of RS lie in the Wingless-related Integration site-1 (WNT) (Table S1). Of these, Receptor tyrosine kinase-like orphan receptor 2 (*ROR2*) and Nucleoredoxin (*NXN)* cause autosomal recessive RS (ARRS) (White et al., 2018; Zhang et al., 2022) while Dishevelled (*DVL1*, *DVL2*, *DVL3)*, *WNT5A*, Frizzled2 (*FZD2*) have autosomal dominant inheritance (ADRS) (Table S1) (Nagasaki et al., 2018; Person et al., 2010; White et al., 2015; White et al., 2018; White et al., 2016; Zhang et al., 2022). The primary phenotypes of RS include facial defects (hypertelorism or wide-spaced eyes, broad forehead, flat nasal bridge)(Beiraghi et al., 2011; Conlon et al., 2021; Kaissi et al., 2020; Sakamoto et al., 2021), limb shortening (Abu-Ghname et al., 2021; Zhang et al., 2022), and genital defects (males, small and buried penis; females, hypoplastic labia majora, small clitoris)(Patton and Afzal, 2002; Roifman et al., 2019). Despite high genetic heterogeneity, RS individuals have similar clinical presentations (Mazzeu et al., 2007) which suggests that ultimately the genes may share a common, indirect, downstream mediator in the WNT signaling pathway. Previously, our lab has investigated effects of ADRS *WNT5A* variants in jaw (Hosseini-Farahabadi et al., 2017) and limb development (Gignac et al., 2019), *DVL1* variants in limb development (Gignac et al., 2023), and here we are focused on two ADRS-*FZD2* variants and how they affect craniofacial development and WNT signaling.

WNTs are secreted glycoproteins that trigger the – (i) canonical or β-catenin-dependent (Nusse and Clevers, 2017; Rim et al., 2022) and (ii) several non-canonical or β-catenin-independent pathways (Angers and Moon, 2009; Lojk and Marc, 2021; Riquelme et al., 2023; Rogers and Scholpp, 2022). Most of the genes implicated in pathogenesis of RS function in the non-canonical c-Jun Planar Cell Polarity (JNK/PCP) pathway expect for *FZD2, DVL1,2,3* that also operate in the canonical/β-catenin-mediated WNT pathway. In the canonical pathway, WNT ligands binds to *FZD* and Low-density lipoprotein receptor-related protein5/6 (*LRP5/6*) receptors recruitment of *DVL* and destabilization of the β-catenin destruction complex. The free β-catenin accumulates in the cytoplasm and then translocated into the nucleus, where it forms a complex with the T-cell factor/Lymphoid Enhancer Factor (TCF/LEF) family of transcription factors and activates transcription of WNT target genes (Nusse and Clevers, 2017). The translocation of β-catenin to the nucleus is a major signaling step in the WNT pathway and is often used as a read out of active canonical pathway (Anthony et al., 2020; Valenta et al., 2012). In the non-canonical JNK/PCP pathway, the WNT ligands bind with either FZD-ROR heterodimer (Green et al., 2014) or ROR homodimers leading to recruitment of DVL which acts as a branch point for two small GTPase (Rac, Rho) pathways (van Amerongen et al., 2012). The Rac-mediated pathway involves activation of c-Jun N-terminal kinase (JNK) and which directly regulates cell polarity (Habas et al., 2003; Topczewski et al., 2011; Widelitz, 2005). Since multiple genes are involved, the pathogenesis of RS could result from an imbalance of either branch of WNT signaling pathways, as we recently reported for the *DVL1* variants (Gignac et al., 2023) and others for reported variants in *FZD2* (Liegel et al., 2023; Zhu et al., 2023).

FZD2 belong to the FZD family of ten receptors that have redundant functions (Schulte, 2010). Our lab and others have shown that FZD2 is expressed abundantly in the developing face (Geetha-Loganathan et al., 2009; Yu et al., 2010; Yu et al., 2012). The extracellular region of all FZDs consists of an N-terminal signal sequence followed by a highly conserved cysteine-rich ligand domain (CRD). The CRD is linked to the seven-pass transmembrane domain consisting of three extra- and three-intracellular loops and a C-terminus. The third intracellular loop and C-terminus are essential for interaction with Dishevelled (Figure S1)(Schulte, 2010). *FZD2* is crucial for embryogenesis, cell polarity, cell proliferation, and many other processes in developing and adult organisms (Huang and Klein, 2004; Wang et al., 2016). *FZD1*, *FZD2,* and *FZD7* share 75% sequence homology and have substantial redundancy (Yu et al., 2010; Yu et al., 2012). *Fzd2^-/-^* null mice have fully penetrant cleft palate (Yu et al., 2010). The craniofacial phenotypes were more severe when double mutants (*Fzd1^-/-^,Fzd2^-/-^, Fzd2^-/-^,Fzd7^-/-^*) were created (Yu et al., 2010; Yu et al., 2012). Recently, *Fzd2^-/-^* null mice was determined not to be a complete null and therefore a new *Fzd2* floxed mouse was generated that had early embryonic lethality (Michalski et al., 2021). Two frameshift mutations in the DVL binding domain of *Fzd2* were created. A single-nucleotide insertion in *Fzd2* (*Fzd2^em1Smill^*) caused cleft palate (CP)(Zhu et al., 2023) and mice with nonsense mutations in *Fzd2 (Fzd2^W553*^)* (Liegel et al., 2023) are born with CP and other craniofacial deformities. This suggests that *Fzd2* plays crucial roles in craniofacial development. In contrast, individuals with *FZD2*-associated ADRS have mild phenotypes (Zhang et al., 2021). Therefore, other animal models that model more closely the RS features are needed.

*FZD2* is the second most common ADRS-causing gene (Zhang et al., 2022). To date, there are 17 ADRS patients with missense or truncating variants in *FZD2* (Table S2). Here, we investigate two missense *FZD2* variants using the chicken embryo model. The missense *FZD2* variants (c.*425C>T* codes for p.Pro142Leu, c.*1301G>T*, codes for p.Gly434Val) were chosen based on their location, unclear pathogenicity, and facial phenotypes. The *425C>T*, termed as a variant of uncertain significance (VUS), lies in the CRD (Fig. S1A,B,B’). *425C>T* was identified in one compound heterozygous individual (*425C>T*, 1130G>A) (White et al., 2018). The father and two siblings of the proband also carry the *425C>T* variant, have short stature but no facial defects. The mother of the proband carries truncating *FZD2* variant (*1130G>A*; Trp377*) and has milder form of ADRS (short stature, broad forehead) (White et al., 2018). The second variant we characterized (*1301G >T*) was identified in the majority of ADRS patients and correlates with more severe face and limb phenotypes. ADRS patients carry *FZD2* variants that alter the Glycine 434 to either serine or valine (Nagasaki et al., 2018; Saal et al., 2015; Türkmen et al., 2017; Warren et al., 2018; White et al., 2018; Zhang et al., 2022). Glycine 434 present in the third intracellular loop of *FZD2* is highly conserved (Fig. S1A,C,D), and is essential for DVL binding (Tauriello et al., 2012). The elevated occurrence of pathogenic variants at Glycine 434 can induce steric hindrance, leading to a reduced FZD-DVL affinity or stability (Punchihewa et al., 2009; Türkmen et al., 2017) (Fig. S1C,D). The *1301G>T* variant was associated with pathogenesis of autosomal dominant Omodysplasia type II (OMODII)(Nagasaki et al., 2018; Saal et al., 2015; Türkmen et al., 2017; Warren et al., 2018; Zhang et al., 2021). Given the significant overlap in phenotypes between OMODII and ADRS, the OMODII is currently recognized as part of ADRS caused by *FZD2* variants (Zhang et al., 2022).

We use the chicken embryo which is a time and cost-effective animal model to study missense variants associated with dominant disease. Our approach involved introducing avian-specific replication competent retroviruses containing wt or variant human *FZD2* into the frontonasal mass, a facial prominence most affected in RS patients (Fig. 1A). The frontonasal mass in birds forms the prenasal cartilage and the flanking premaxillary intramembranous bones (Abramyan and Richman, 2018). In humans, the frontonasal mass contributes to midline structures of the face encompassing the bridge of the nose, the central region of the nose, the nasal septum, the philtrum, and the central portion of the upper lip, including the premaxilla (Chen et al., 2013). Previous studies from our lab have demonstrated that ADRS-*WNT5A* (Gignac et al., 2019; Hosseini-Farahabadi et al., 2017) and *DVL1* variants (Gignac et al., 2023) have dominant-negative effects on chondrogenesis leading to abnormal bone morphology mimicking ADRS. In this study, we uncover functional evidence that both *425C>T* and *1301G>T* weakly activate the canonical WNT pathway, have dominant negative effects on activity of wt *FZD2* and inhibit skeletogenesis.

**Figure 1:**
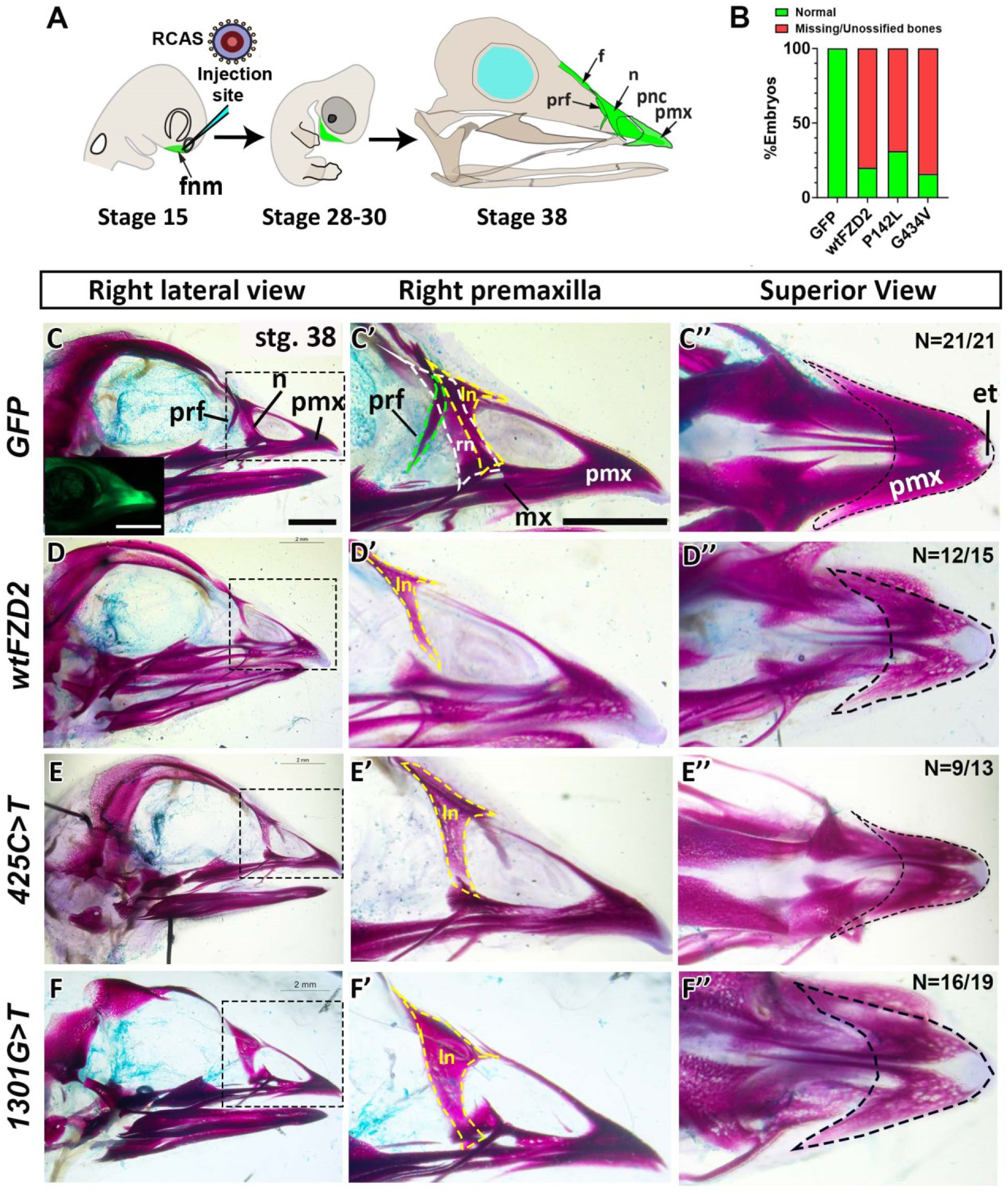
In vivo overexpression of wt and variant h*FZD2* viruses inhibit upper beak patterning and ossification. (A) Illustration of microinjection of RCAS viruses into the frontonasal mass at stage 15 (E2.5), analysis at stage 28 (E5.5) and stage 38 (E12.5). B) Fraction of total analysis of normal and abnormal embryos (missing/unossified bones). (Inset in C) An embryo injected with *GFP* showing viral spread as green fluorescence in the upper beak. (C-F’’) Right lateral view of skulls stained with Alcian blue and Alizarin red (stage 38) injected with *GFP* (n=21), wt (n=15) or h*FZD2* variants (*425C>T* n=13, *1301G>T* n=19). (C’) Magnified right lateral view of *GFP*-injected embryos with normal prefrontal (green dash line), right nasal (white dash line) and left nasal (yellow dash line) bones. (D’,E’,F’) Missing prefrontal right and nasal bones in embryos injected with h*FZD2* viruses, left nasal bone was present. (C’’) Magnified superior view of *GFP* embryos showing alizarin red stained premaxillary bone (black dash line). (D’’,E’’,F’’) Comparatively faint stain was seen in h*FZD2 I* viruses. The reduced Alcian blue stain is a technical error. Key: et; egg tooth, f; frontal, fnm; frontonasal mass, ln, left nasal; mx, maxilla, na; nasal, prf, prefrontal; pmx, premaxilla; pnc; prenasal cartilage, rn, right nasal. Scale bar: C-F’’= 2mm.

## RESULTS

In these experiments we compare the effects of generalized increased levels of wild-type (wt) h*FZD2* to the *GFP* (control) and two ADRS h*FZD2* variants (*425C>T* and *1301G>T*). The *GFP* containing virus is suitable control for overexpression studies since it does not affect development (Geetha-Loganathan et al., 2014; Hosseini-Farahabadi et al., 2017). The strategy used here is to express the human genes (wt or variant h*FZD2*) in defined regions of the chicken embryo. Since the ADRS patients carry one normal and one abnormal copy of *FZD2* gene, overexpression of variant *FZD2* alongside the endogenous chicken genome (containing g*FZD2*) is appropriate for studying autosomal dominant mutations. Indeed, our previous work on autosomal dominant *DVL1* frameshift mutations (*1519*Δ*T, 1529*Δ*G, 1615*Δ*A*) (Gignac et al., 2023) and *WNT5A* missense mutation (*248G>*C) (Gignac et al. 2019; Hosseini-Farahabadi et al. 2017) found that the variant *WNT5A* was sufficient to reduce size and change bone shape. Here, the data will be interpreted as follows: (1) If the proteins coded by missense h*FZD2* variants expressed on top of the Gallus *FZD2* lacked function, then treated embryos should look like *GFP* controls, (2) If the variants caused no additional effects other than those caused by overexpression of h*FZD2*, then phenotypes are due to generally higher levels of h*FZD2* rather than specific effects of the variants, (3) If the phenotypes were similar but more severe than wt h*FZD2*, then the variants caused a gain-of-function, and finally (4) if the variant viruses induced de novo phenotypes compared to wt h*FZD2*, then this would suggest more complex functional alterations that would require additional analysis.

### Overexpression of h*FZD2* in vivo leads to abnormal patterning and inhibition of ossification

To determine the skeletal phenotype induced by the exogenous h*FZD2*, the embryos injected at stage 15 [embryonic day (E) 2.5] (Hamburger and Hamilton, 1992) were incubated until stage 38 [E12.5], allowing analysis of 3D skeletal phenotypes. Examination of heads showed that most of the embryos infected *GFP* (control), wt h*FZD2* or variant h*FZD2* had normal beak morphology (Table S3) and survival ranged between 64-78% (Table S4). In cleared skeletal preparations (stage 38, 10 days post-injection) stained with Alcian blue (to detect cartilage) and Alizarin red (to detect mineralized bone), *GFP* infected skulls had all frontonasal mass derivatives (premaxilla, nasal, prefrontal bones) with robust alizarin red staining (Fig. 1C-C’’; Fig. S2A-C). The spread of the *GFP* virus in the upper beak confirmed accurate targeting (Fig. 1C’ inset). Decreased bone staining and agenesis of bones were observed in most embryos injected with wt h*FZD2* or variant h*FZD2* (Fig. 1B, D-F’’, Fig. S2D-L). In particular, the prefrontal and the nasal bones were missing while the premaxillary bone showed faint or no alizarin red staining compared to *GFP* (Fig. 1D-F’’; Fig. S2A-L; Fig. S3A-C, Table S3). The absence of frontonasal derivatives and reduced ossification in vivo suggested that having generally higher levels of exogenous h*FZD2* protein, whether normal or variant, inhibits intramembranous bone formation in chicken embryos.

### Overexpression of variant h*FZD2* viruses caused increase in frontonasal mass width mimicking ADRS

To understand the molecular mechanism underlying the skeletal phenotypes we examined embryos prior to the onset of cell differentiation, at stage 28 (E5.5). We sectioned the embryos and used an antibody that recognizes the viral structural protein GAG, to confirm that virus was localized in the frontonasal mass (Fig. 2 A-D, Fig. S4A-X). Next, we selected embryos with strong GAG expression in the frontonasal mass for further molecular characterization (Fig. S4A-X). Due to the virus being replication competent and the inability to quantify the initial injected virus amount, there was variability in viral spread. Serial sections were stained for the early chondrogenic marker SOX9 (SRY-Box Transcription Factor 9) and TWIST1 (Twist-related protein 1), a marker for undifferentiated mesenchymal cells. The wt and variant viruses expressing FZD2 protein, did not affect the differentiation of SOX9-positive prenasal cartilage (Fig. 2E-H). The TWIST1-positive undifferentiated mesenchyme was also unaffected (Fig. 2I-L). We also analyzed cell proliferation (BrdU) and apoptosis (TUNEL) to assess any potential reduction in cells contributing to the formation of the beak skeleton. There were no differences in cell proliferation (Fig. 2M-P; V, Table S5) or cell death (Fig. 2Q-T’, W, Table S5) compared to *GFP*. Since ADRS individuals have wider faces, we analyzed the width of the frontonasal mass in the presence of wt or variant viruses (schematic in Fig. 2X). Interestingly, all embryos infected with variant h*FZD2* viruses had wider frontonasal mass compared to wt h*FZD2* and *GFP* infected embryos (Fig. 2Y). This phenotype closely recapitulates the broad face phenotypes observed in ADRS individuals.

**Figure 2:**
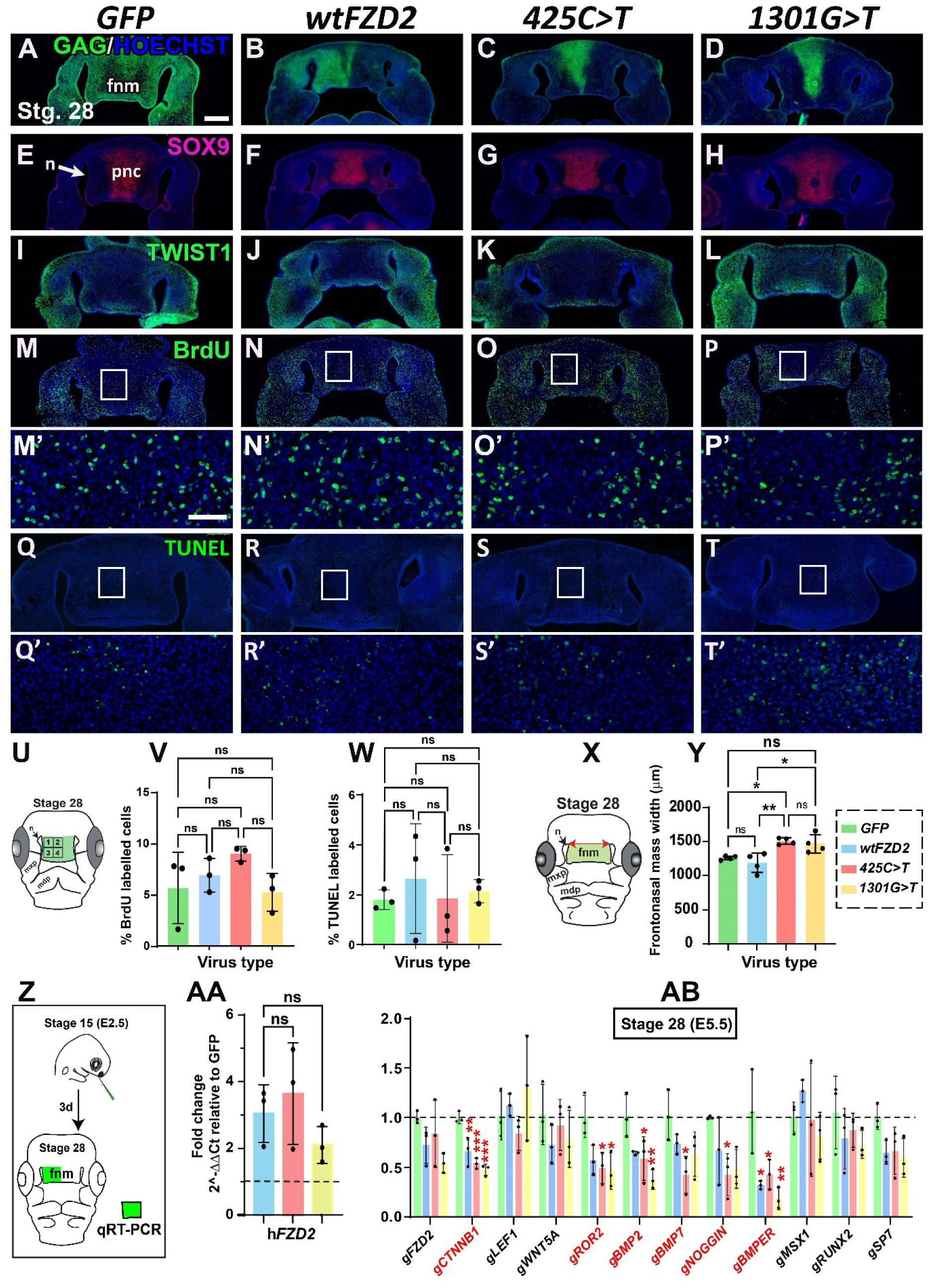
Immunostaining on stage 28 embryos injected with *GFP*, wt h*FZD2* and h*FZD2* variants. Frontal sections of heads injected at stage 15 and fixed 72 h post injection at stage 28 (A-D) GAG-staining shows viral spread in the frontonasal mass (n=7). (E-H) Serial sections show SOX9 in prenasal cartilage and (I-L) expression of undifferentiated mesenchymal marker TWIST1 surrounding SOX9-positive area. BrdU was injected 2 h prior to euthanasia. (M-P) Near adjacent sections stained with anti-BrdU (n=3) and (M’-P) magnification of white boxes in the frontonasal mass. (Q-T) TUNEL assay performed on a different set of embryos (n=3) and (Q’-T’) magnification of white boxes in Q-T. Quantification of (V) BrdU and (W) TUNEL positive cells in the frontonasal mass show no differences between viruses. (X) Schematic showing measurement of frontonasal mass width (n=4). (Y) The variant h*FZD2* embryos had wider frontonasal mass compared to wt h*FZD2* and *GFP*. (Z) Schematic for collection of frontonasal mass for qRT-PCR analyses. Levels of expression were compared to *GFP* using ΔΔCt. (AA) The expression of h*FZD2* was 2-5-fold higher than *GFP*. (AB) RNA expression profile showing effects of h*FZD2* viruses on WNT and BMP pathway genes. The statistical analysis was done with one-way ANOVA, Dunnett’s test in GraphPad Prism 10.1.0. The error bars represent one standard deviation. Key: fnm, frontonasal mass; mxp, maxillary prominence, mdp; mandibular prominence, n - nasal slit, ns- not significant; pnc, prenasal cartilage, stg; stage, p*<0.05, **p<0.01. Scale bar in A-P = 500µm, Q-T = 200µm, Q’T’ and U’-X’ = 20µm, U-X= 100µm

### The h*FZD2* viruses inhibit ossification by synergistic downregulation of WNT and BMP pathways

The effects of the viruses on gene expression were quantified using qRT-PCR. Notably, the addition of exogenous h*FZD2* gene did not affect expression of endogenous *Gallus FZD2* (g*FZD2*) (Fig. 2Z). Since WNT and Bone Morphogenic Protein (BMP) pathway mediate osteogenesis independently and synergistically (Itasaki and Hoppler, 2010; Lin and Hankenson, 2011), we measured WNT and BMP pathway mediators (Fig. 2Z). In wt and variant h*FZD2* frontonasal mass tissues, the transcript levels of g*CTNNB1* (canonical pathway mediator) and g*ROR2* (non-canonical WNT receptor) were significantly downregulated compared to *GFP-* infected tissues (Fig. 2AB). This suggested that the h*FZD2* viruses cause impairment of both canonical and non-canonical WNT pathways. Loss-of-function variants in *ROR2* cause autosomal recessive RS (Table S1). The indirect downregulation of g*ROR2* by exogenous variant h*FZD2* viruses may explain the similar but mild clinical phenotypes caused by the h*FZD2* variants.

In the BMP pathway, the *425C>T* variant significantly downregulated *BMP2*, *BMP7*, *NOGGIN*, and *BMPER,* while *1301G>T* variant downregulated *BMP2* and *BMPER* (Fig. 2AB). The decrease in *BMPER* by wt and variant h*FZD2* viruses is intriguing since the BMPER protein can either increase or decrease BMP signaling, depending on the context (Correns et al., 2021). There was no change in expression of g*LEF1*, g*WNT5A* at stage 28 and g*MSX1*, g*RUNX2*, g*SP7* (Fig. 2.AB). There were no differences in the expression of WNT and BMP pathway mediators between the wt and variant h*FZD2* viruses (Fig. 2AB). Overall, the RNA expression prolife at early stages of embryonic development implies that the wt and variant h*FZD2* viruses synergistically downregulated WNT (g*CTNNB1*) and BMP (g*BMP2*, g*BMP7*, g*NOGGIN*, g*BMPER*) pathways. The synergistic crosstalk between the WNT and BMP pathways, coupled with prolonged suppression of BMP regulator, *BMPER*, may have ultimately contributed to inhibition of upper beak patterning and ossification caused by all FZD2 viruses (Itasaki and Hoppler, 2010; Xiao et al., 2018).

### Variant *FZD2* viruses inhibit chondrogenesis in frontonasal mass micromass cultures

Due to the observed downregulation of WNT and BMP pathways in vivo, crucial pathways that regulate skeletogenesis, we characterized the h*FZD2* variants using high-density frontonasal mass micromass cultures. Micromass cultures are a well-established approach to study chondrogenesis in the face (Hosseini-Farahabadi et al., 2013; Richman and Crosby, 1990) and limb (Weston et al., 2002). Viral spread was observed as strong green fluorescence on day 4, 6 and 8 (Fig. 3A,J,S). All virus infected cultures were stained in wholemount with Alcian blue (to detect cartilage sheets) and alkaline phosphatase (to detect mineralized chondrocytes, day 6 and 8) (Fig. 3AB). On day 4, all viruses allowed formation of Alcian blue stained cartilage condensations observed in wholemount stained cultures (Fig. 3B-E,AB) and in histological sections (Fig. 3F-I). This implies that the wt and variant h*FZD2* viruses have no effect on the early stages of chondrogenesis, such as differentiation of primary face mesenchyme into chondrocyte lineage and formation of cartilage condensations. At day 6, frontonasal mass formed robust Alcian blue-positive cartilage sheets (Fig. 3K-N). Alkaline phosphatase-positive mineralized chondrocytes were visible at this stage in wholemount cultures (Fig. 3K-N,AB). On day 8, mineralization had progressed (Fig. 3T-W’). Quantitative differences in Alcian blue or alkaline phosphatase staining could not be accurately determined from wholemount images. Transverse histological sections allowed measurements of cartilage matrix. Another reason we used sections for the majority of analyses is that the presence of viral protein within the cultures could confirmed using immunostaining (Fig. 4E-H,Q-T; sample sizes Table S7)

By day 6, thick cartilage sheets formed in the center of the cultures infected with GFP, wt and the *425C>T* variant h*FZD2* viruses (Fig. 3O-R). Interestingly, cultures infected with the variant (*1301G>T*) h*FZD2* viruses were significantly thinner compared to all other viruses (Fig. 3O-R, AC). By day 8, cultures treated with wt or variant h*FZD2* viruses were significantly thinner compared to *GFP* infected cultures (Fig. 3X-AA; AD). Moreover, quantification of area occupied by Alcian blue in sections showed that the variant h*FZD2* viruses significantly inhibited chondrogenesis compared to *GFP* treated cultures (Fig. 3AE). The *425C>T* cultures had a significantly lower amount of cartilage compared to the *1301G>T* (Fig. 3AE), the first time a difference between the variants was detected. Overall, the data suggests that the variants of *FZD2* affect later stages of chondrocyte differentiation and have inhibitory effects on chondrogenesis *in vitro*.

**Figure 3:**
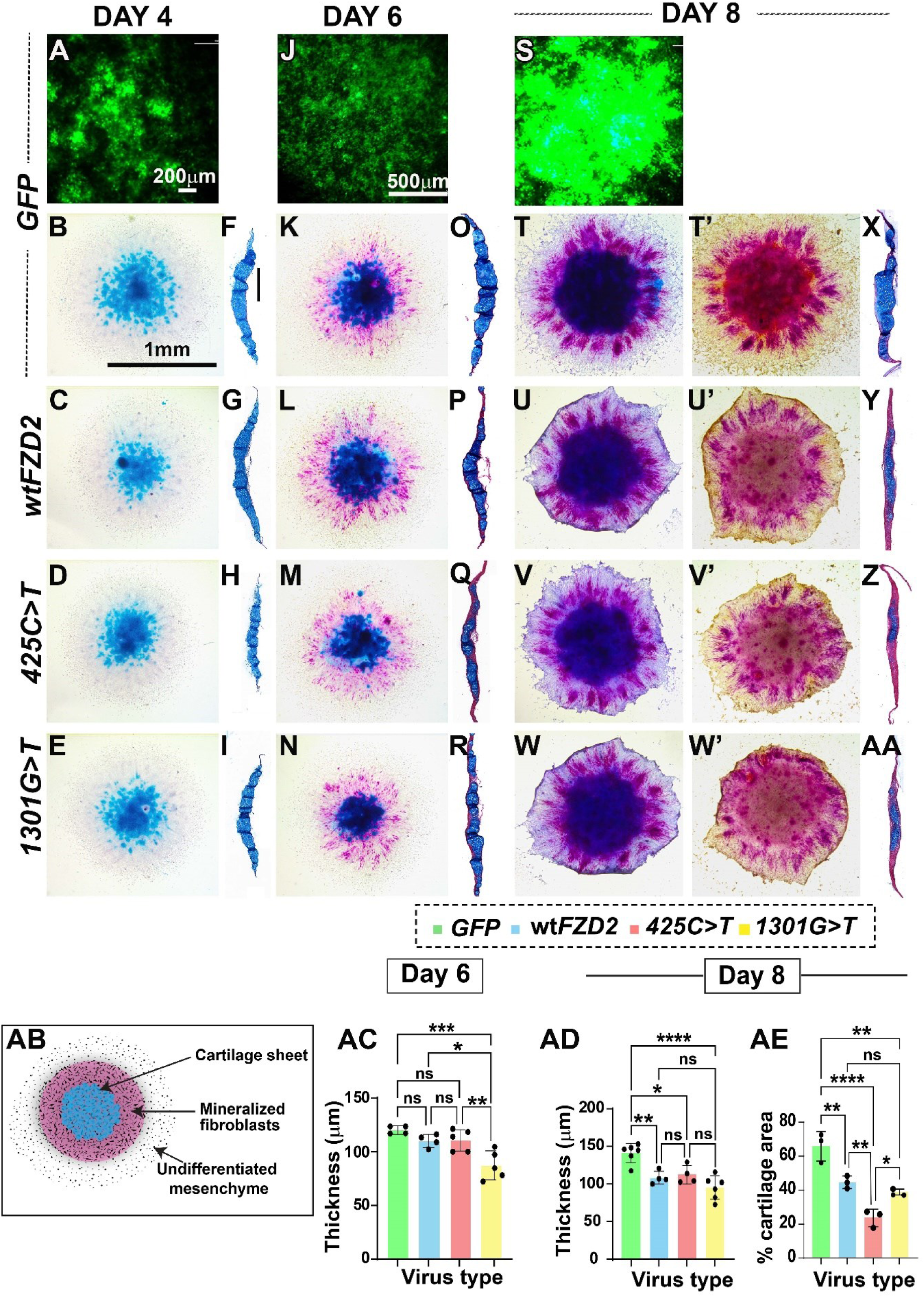
Wholemount and histological analysis of day 4, 6, and 8 cultures. *GFP* virus spread in frontonasal mass cultures observed on days 4 (A), 6 (J), and 8 (S). (B-E) Day 4 wholemount cultures were stained with Alcian blue for cartilage detection and counterstained with hematoxylin, showing normal cartilage (F-I) Transverse sections on day 4 (n=3)also displayed normal cartilage. (K-N) Wholemount day 6 cultures stained with Alcian blue and alkaline phosphatase (to detect mineralized chondrocytes). (O-R) day 6 (n=7-9) histological sections stained with Alcian blue and Picrosirius red. (T-W’) By day 8, h*FZD2* and *GFP* treated cultures stained with Alcian blue and alkaline phosphatase showed no differences, although h*FZD2* cultures had less alkaline phosphatase staining. (X-AA) h*FZD2* cultures remained thinner than *GFP*. (AB) Illustration showing wholemount culture areas. (AC) Day 6 histological measurements indicated *1301G>T* cultures were thinner than others (AD) By day 8 (n=4-6), all h*FZD2* mutant cultures were significantly thinner than *GFP*. (AE) with all h*FZD2* viruses showing significantly lower cartilage area. The *425C>T* virus notably inhibited chondrogenesis more than wild-type h*FZD2* and *1301G>T*. The statistical analysis was done with one-way ANOVA, Dunnett’s test in GraphPad Prism 10.1.0. The error bars represent one standard deviation. Scale bar in A=200μm, scale bar in B=1mm applies to C-E, K-N, T-W’, scale bar in F = 100μm applies to G-I, O-R, X-AA, J and S = 500μm. p values: *≤0.05, **≤0.01, ***≤0.001, ****≤0.0001, ns – not significant.

**Figure 4:**
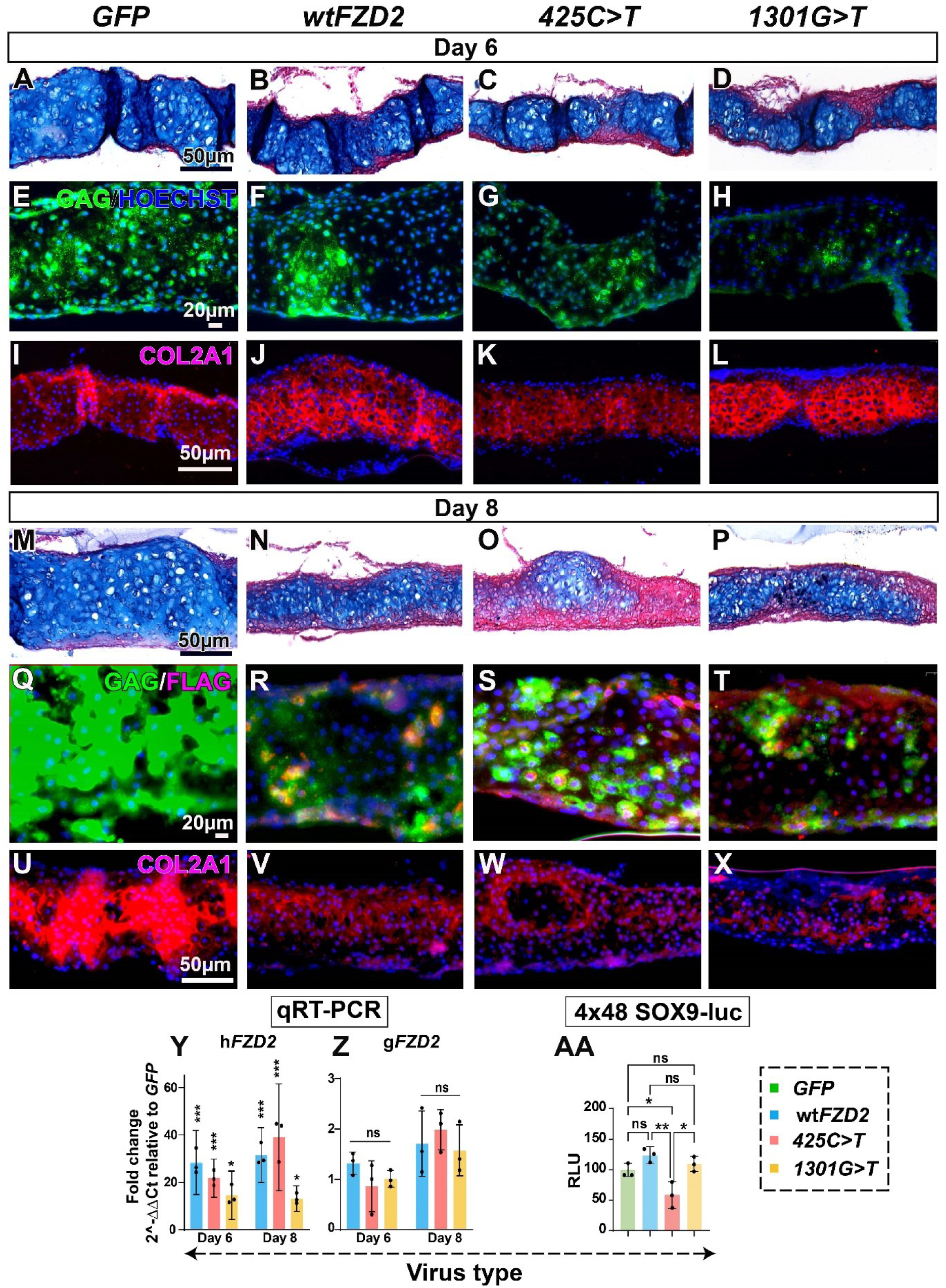
Effects of h*FZD2* viruses on cartilage matrix in vitro on day 6 and day 8. (A-D) Magnified view of day 6 cultures stained with Alcian blue and picrosirius red. (E-H) Near adjacent sections GAG-positive sections showing viral spread (n=7-9). (I-L) Immunostaining of serial sections show comparable amounts of cartilage matrix protein (type II collagen;COL2A1). (M-P) Histological sections of day 8 cultures (n=4-6) showed reduced Alcian blue and thickness in h*FZD2* infected cultures compared to *GFP*. (Q-T) Viral spread shown with anti-GAG and anti-FLAG staining. (U-X) All h*FZD2* infected cultures show weak COL2A1 expression compared to *GFP*. A separate set of cultures was collected for qRT-PCR analysis on day 6 and day 8. Levels of expression were compared to *GFP* using ΔΔCt. (Y) In h*FZD2* infected cultures h*FZD2* transcript was 15-to-40-fold higher compared to *GFP* cultures while (Z) the levels of g*FZD2* were unchanged. The statistical analysis was done in GraphPad Prism 10.1.0 with one-way ANOVA, Dunnett’s test. (AA) Micromass cultures were transfected with SOX9 luciferase reporter 24 h after plating and reading was done on day 3. Cultures infected with *425C>T* had significantly reduced SOX9 activity compared to other viruses. The statistical analysis was done with one-way ANOVA, Tukey’s post hoc test in GraphPad Prism 10.1.0. All error bars represent standard deviation. Scale bar in A= 50µm applies to B-D, M-P, U-X Scale bar in E= 20µm applies to F-H, Q-T Scale bar in I = 50µm applies to J-L. Key: p*<0.05, ** p<0.01, ***p<0.001, ns – not significant

All day 6 and 8 cultures showed comparable viral spread (Fig. 4E-H, Q-T). On day 6, robust type II collagen (COL2A1) expression was observed, consistent with the Alcian blue staining observed in histological sections (Fig. 4A-D,I-L). There was no discernible difference between the wt h*FZD2*, h*FZD2* variants and *GFP* (Fig. 4I-L). On day 8, all h*FZD2*-infected cultures showed weak COL2A1 staining compared to *GFP* (Fig. 4U-X). This data is consistent with reduced Alcian blue staining observed in histological sections (Fig. 4M-P,U-X).

A 15-40-fold increase in expression of exogenous h*FZD2* (Fig. 4Y) did not alter the expression endogenous g*FZD2* (Fig. 4Z) showing there were no feedback loops induced. However, on day 6, g*CTNNB1* was downregulated by both variant h*FZD2* variants while g*WNT5A* was downregulated by the wt h*FZD2* (Fig. S5A). By day 8, no differences in gene expression were detected (Fig. S5B) suggesting that homeostasis was restored, even though the cartilage was significantly thinner for the FZD2-infected cultures.

To further explore the mechanism for the reduction in cartilage, we used a reporter that measures the level of SOX9 transcriptional activity (Weston et al., 2002). *SOX9* has been shown to directly regulate *COL2A1* in chondrocytes (Bell et al., 1997; Ng et al., 1997). The reporter was transfected 1 day after plating to time the biochemical readout of luciferase activity to day 3 of the culture. The cartilage condensations are first visible at 48h post plating (Hosseini-Farahabadi et al., 2013). Interestingly, the *425C>T* virus failed to activate the SOX9-luciferase reporter (Fig. 4AA). This suggests that at least for this variant, lower SOX9 expression could have reduced the number of chondrocytes contributing to lower cartilage matrix deposition.

We next analyzed whether h*FZD2* viruses reduced cell proliferation and increased cell death thereby contributing to reduced thickness of the cultures. On day 6, the BrdU labelled proliferating chondrocytes throughout the cartilage forming area (Fig. S6A-D; n = 7-9, Table S7). However, by day 8, the there were almost no proliferating cells in the culture (Fig. S6E-H, n = 4-6, Table S7). There was no visible difference in cell proliferation between the viruses. On day 6, there were fewer apoptotic cells compared to day 8 across all virus infected culture conditions (Fig. S6I-P). Quantification revealed no effects of the wt or variant h*FZD2* on apoptosis (Fig. S6Q, R). Overall, data indicates that the thinning of cultures is not related to reduction of proliferation or increased apoptosis but is due to deficiency in extracellular matrix synthesis.

### Prolonged TWIST expression is seen in variant *FZD2*-infected cultures

To investigate the possible role of canonical WNT signaling in mediating the chondrogenic phenotypes of h*FZD2* infected cultures, we examined expression of nuclear β-catenin, a marker of active canonical WNT signaling. In *GFP* and wt h*FZD2* infected cultures, most nuclear β-catenin expressing cells were found at the periphery of the cultures and excluded from the cartilage (Fig. 5A, B). In contrast, in *425C>T* and *1301G>T*-infected cultures, chondrocytes residing in the cartilage matrix also expressed nuclear β-catenin (Fig. 5C, D). Quantification of the proportion of cells expressing nuclear β-catenin revealed that *425C>T* and *1301G>T* infected cultures had significantly higher number of nuclear β-catenin positive chondrocytes compared to *GFP* and wt h*FZD2* (Fig. 5I). This implies that the variants maintain canonical β-catenin mediated signaling longer than wtFZD2. We attempted to measure the levels of *CTNNB1* RNA in day 8 cultures but there was no significant difference (Fig. S5B). The mechanism for decreased cartilage in wtFZD2 appears to be different than the variants and does not involve translocation of β-catenin to the nucleus.

**Figure 5:**
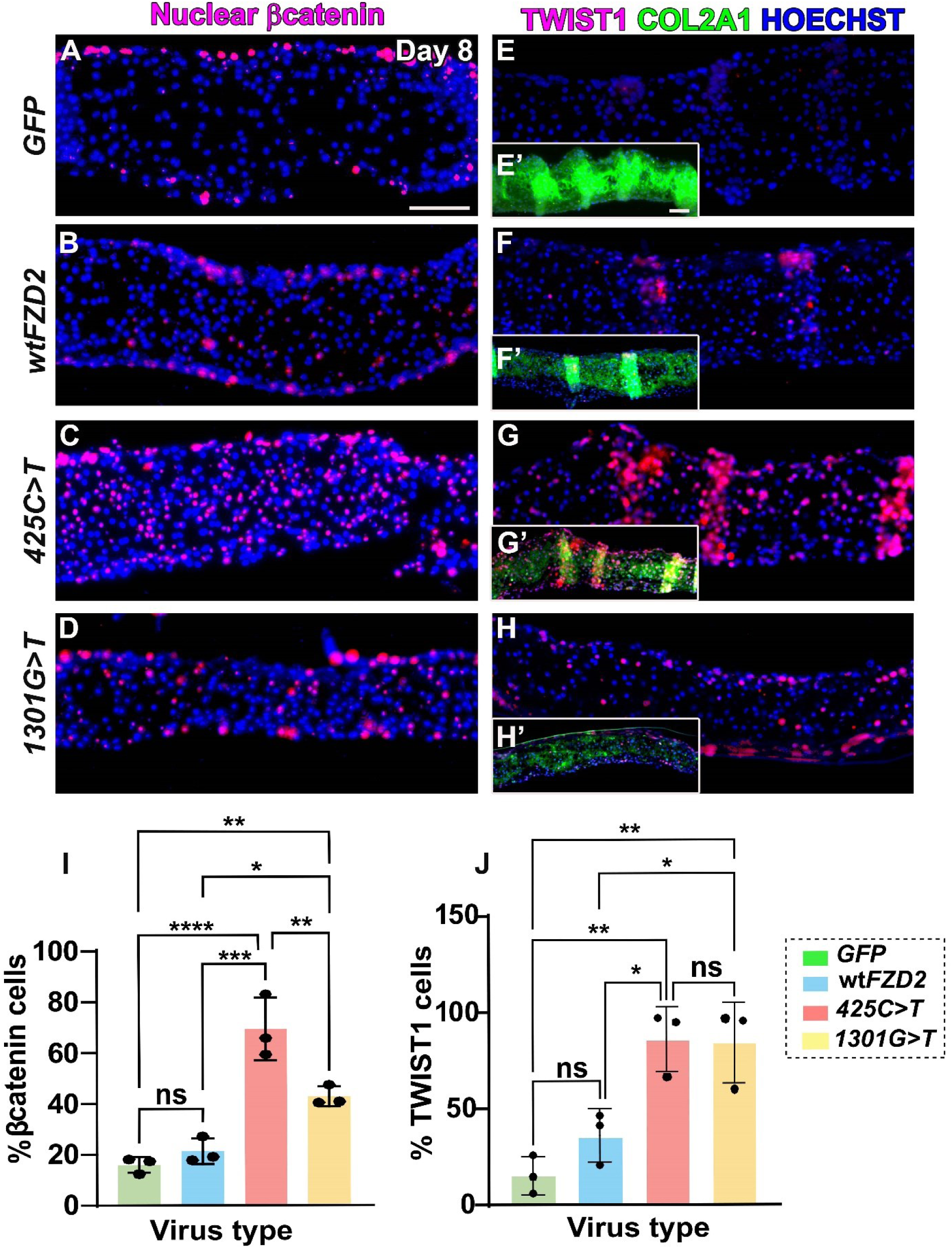
h*FZD2* variants increased expression of nuclear β-catenin and TWIST1 on day 8. (A, B) In *GFP* or wt h*FZD2* virus infected cultures, nuclear-β-catenin was found in fibroblasts at the culture edges. (C, D) Conversely, in variant hFZD2 cultures, nuclear-β-catenin showed high expression within cells in cartilage-forming regions. (E-H) Serial sections showing expression of TWIST1 and (E’-H’) show colocalization of TWIST1 and COL2A1. (E-F) *GFP* and wt h*FZD2* cultures express low levels of TWIST1 while (G, H) variant h*FZD2* viruses have higher proportion of TWIST1-positive chondrocytes. (I) Quantification of proportion of cells expressing nuclear-β-catenin. 40-60% cells in the variant h*FZD2* variant infected cultures expressed nuclear β-catenin. (K) The h*FZD2* variant infected cultures have significantly higher (60-80%) TWIST1-positive nuclei compared to *GFP* and wt h*FZD2*. The statistical analysis was done with one-way ANOVA, Tukey’s post hoc test in GraphPad Prism 10.1.0. The error bars represent one standard deviation. Scale bar = 50µm. Key: p*<0.05, ** p<0.01, ***p<0.001, ns – not significant.

β-catenin (*Ctnnb1*) has been shown to bind with the 5′ enhancer element upstream of the promoter of *Twist1* in cranial mesenchyme (Goodnough et al., 2012). Therefore, as another readout of the β-catenin pathway, we examined if chondrocytes expressed TWIST1, an established repressor of *SOX9* (Goodnough et al., 2012). Interestingly, we observed that the TWIST1-positive chondrocytes were more abundant in the *425C>T* and *1301G>T* variant infected cultures compared to *GFP* (Fig. 5E-H’, J). This suggests that the variant h*FZD2* variants cause β-catenin to enter the nucleus which in turn may activate expression of the chondrogenic repressor TWIST1, ultimately inhibiting chondrogenesis in vitro.

### *FZD2* variants have dominant-negative effects on wt*FZD2* function

Thus far our data suggested the involvement of both canonical and non-canonical WNT pathways in pathogenesis of *FZD2*-associated ADRS. To clarify the activity levels of canonical and non-canonical WNT signaling, plasmids containing either wt or variant h*FZD2* were transiently transfected into frontonasal mass micromass cultures and luciferase assays were carried out. This approach ensured consistency in DNA levels and cell context. Luciferase assays were also conducted using HEK293T cells, a cell line commonly employed for luciferase assays. The Super TOPFlash reporter (STF) is a highly sensitive reporter that detects the βcatenin/TCF-driven transcriptional activity (Biechele and Moon, 2008). In frontonasal mass cultures the addition of wt h*FZD2* strongly activated STF compared to empty vector (pcDNA3.2) (Fig. 6A). This suggested that h*FZD2* activated canonical pathway by utilizing endogenous ligands present in chicken facial mesenchyme. In contrast, STF activation in the presence of the *425C>T* variant was significantly weaker as compared to wt h*FZD2* (Fig. 6A). In HEK293T cells, wt h*FZD2* induced STF activity while both h*FZD2* variants failed to activate the reporter (Fig. 6C). Mouse Fzd2 has been shown to be essential for Wnt3a-dependent accumulation of β-catenin and Wnt5a competes with Wnt3a for binding to Fzd2 (Sato et al., 2010). To test the effects of the *FZD2* variants on ligand interaction, WNT3A was added to the media (100ng/ml) protein. The addition of WNT3A strongly activated STF and this level was significantly elevated in the presence of all h*FZD2* plasmids compared to the empty vector (Fig. 6B). When WNT3A was added to HEK293 cells, there was similar activation of the pathway, regardless of which plasmid was transfected (Fig. 6D).

**Figure 6:**
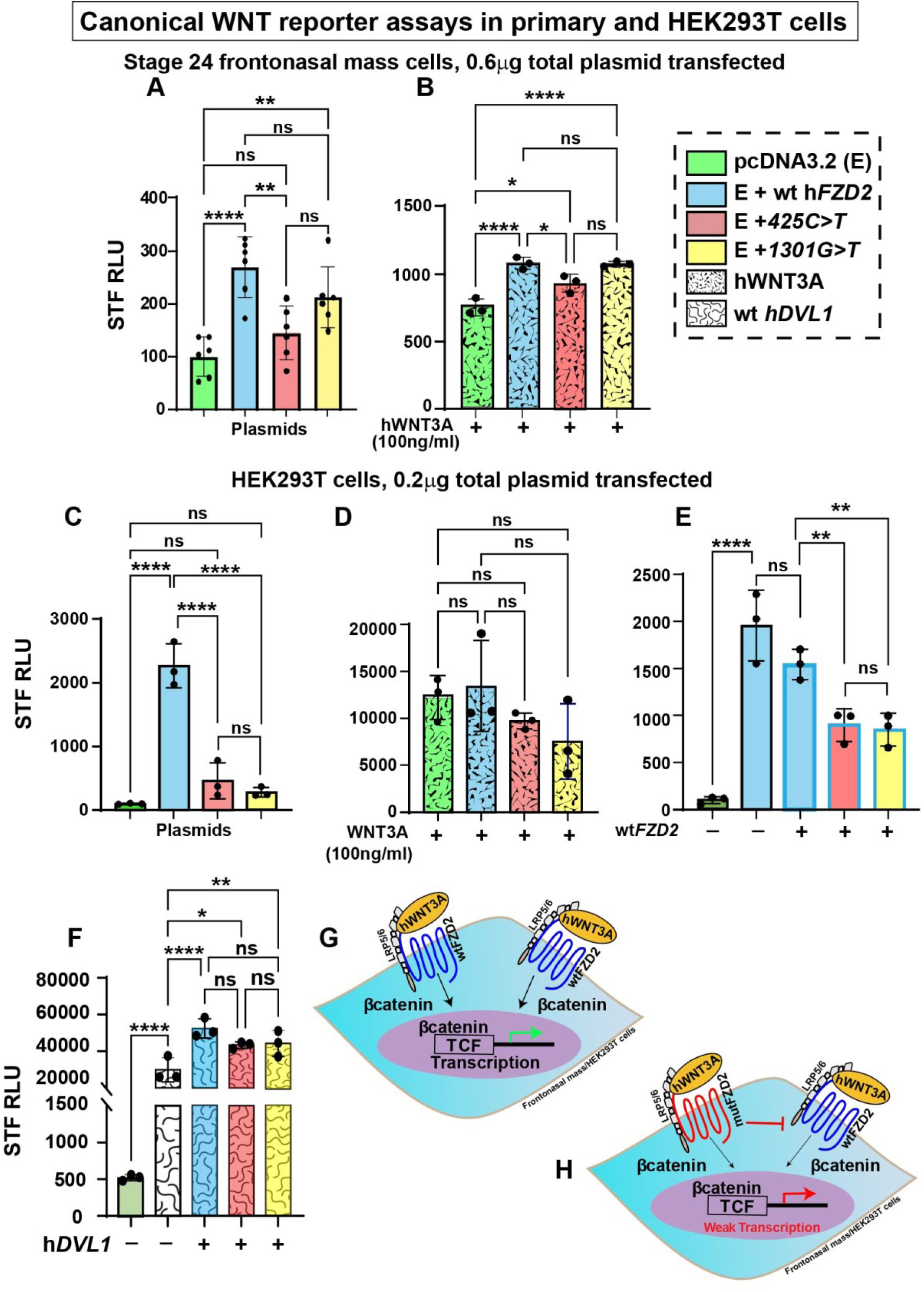
Effects of h*FZD2* variants of WNT canonical signaling pathway in frontonasal mass cultures and HEK293T cells. β-catenin mediated transcriptional activity measured in stage 24 frontonasal mass micromass cultures or HEK293T cells using Super Topflash (STF; M50 Super 8x Topflash) reporter. (A,C) h*FZD2* variants to showed significantly lower activation of STF compared to wt h*FZD2*. (B,D) Addition of recombinant human WNT3A (100ng/ml) 24h post transfection, STF activity increased for all h*FZD2* plasmids. (E) In HEK293T cells, variants combined with equimolar amounts of wt h*FZD2* (0.1ug wt h*FZD2* + 0.1ug empty/wt h*FZD2*/variant h*FZD2*) showed activation of STF compared to wt h*FZD2*. (F) When combined with wt h*DVL1*, all h*FZD2* plasmids showed equally increased STF activity that was significantly higher than h*DVL1* alone. (G-H) Schematic showing activity of wt *FZD2* and RS-h*FZD2* variants. Three biological replicates, three technical replicates and the experiment was repeated three times. Statistical analysis was performed using One-way ANOVA, Tukey’s test. The error bars represent standard deviation. Key: For A and B, 0.6µg of plasmid (0.3 µg of pcDNA3.2 + 0.3µg of *FZD2*) was used for transfection of frontonasal mass cultures; For C-F, 0.2µg of plasmid (0.1 µg of pcDNA3.2/ h*FZD2* + 0.1µg of h*DVL1*) was used for transfection of HEK293T cells; RLU – relative luciferase activity, p values: *≤0.05, **≤0.01, ***≤0.001, ****≤0.0001, ns – not significant.

Since the primary mesenchymal cells showed similar readouts to HEK293T cells, the remainder of the STF luciferase assays were carried out in HEK293T cells. Since FZD receptors form multimers in the cell membrane (Carron et al., 2003; Dann et al., 2001), we next added equimolar amounts of wt h*FZD2* with each of the variants to mimic the heterozygous genotype (autosomal dominant). When either variant was added to wt h*FZD2* there was significantly reduced activation of STF compared to wt h*FZD2* suggestive of a dominant-negative effects (Fig. 6E, illustrated in G, H). This finding is consistent with a study that found that a different RS-associated h*FZD2* variant (*Fzd2^W548*^*) decreases canonical WNT signaling as measured by a reduction in *Axin2* RNA levels (Liegel et al., 2023).

Indeed, the ADRS-*FZD2^W548*^* variant was shown to reduce DVL1 recruitment *in vitro* (Saal et al., 2015). *Fzd2^em1Smill^*mice with a frameshift in the Dvl-interaction domain also have reduced canonical WNT signaling (Zhu et al., 2023), probably due to failure of Dvl1 to interact with Fzd2. Since the *1301G>T* variant lies in the DVL-binding domain, we checked for interactions between h*FZD2* and h*DVL1*. On its own, h*DVL1* significantly activated STF compared to the empty vector and the level was further increased when wt *FZD2* or variant forms were added (Fig. 6F)(Gignac et al., 2023). The wt and variant h*FZD2* variants show comparable canonical WNT activity in the presence of *DVL1.* While the result with the extracellular variant 425C>T was expected, the fact that *1301G>T* did not interfere with DVL1 interactions does not match the predictions made in human genetics studies (Türkmen et al., 2017; White et al., 2018) (Fig. S1). Functional tests are necessary to test in silico modeling.

### *425C>T* variant affecting the extracellular FZD2 domain caused a loss-of-function in the JNK/PCP pathway

Since *FZD2* is known to participate in both canonical and non-canonical WNT pathways (Gordon and Nusse, 2006; Komiya and Habas, 2008), we used Activating Transcription Factor 2 (ATF2)-luciferase (Ohkawara and Niehrs, 2011) to measure the c-Jun Planar Cell Polarity (JNK-PCP) pathway activity. The ATF2 reporter was significantly activated by *wt hFZD2* compared to empty vector pcDNA3.2 (Fig. 7A). The *1301G>T* variant activated ATF2 to the same extent as wt h*FZD2* while *425C>T* did not activate the reporter above basal levels (Fig. 7A). WNT5A (100ng/ml), primarily a non-canonical ligand, slightly activated ATF2 (Fig. 7B) but when WNT5A was combined with wt h*FZD2* or *1301G>T* there were 2-5-fold increase in activity (Fig. 7B). In contrast, the *425C>T* variant - which changes proline to lysine at position 142 in cysteine-rich ligand binding domain - failed to activate ATF2 in the presence of WNT5A (Fig. 7B). We replicated these results in HEK293T cells (Fig. 7C, D). This striking result shows that the *FZD2^425C>T^* variant is unable to utilize the exogenous hWNT5A.

**Figure 7:**
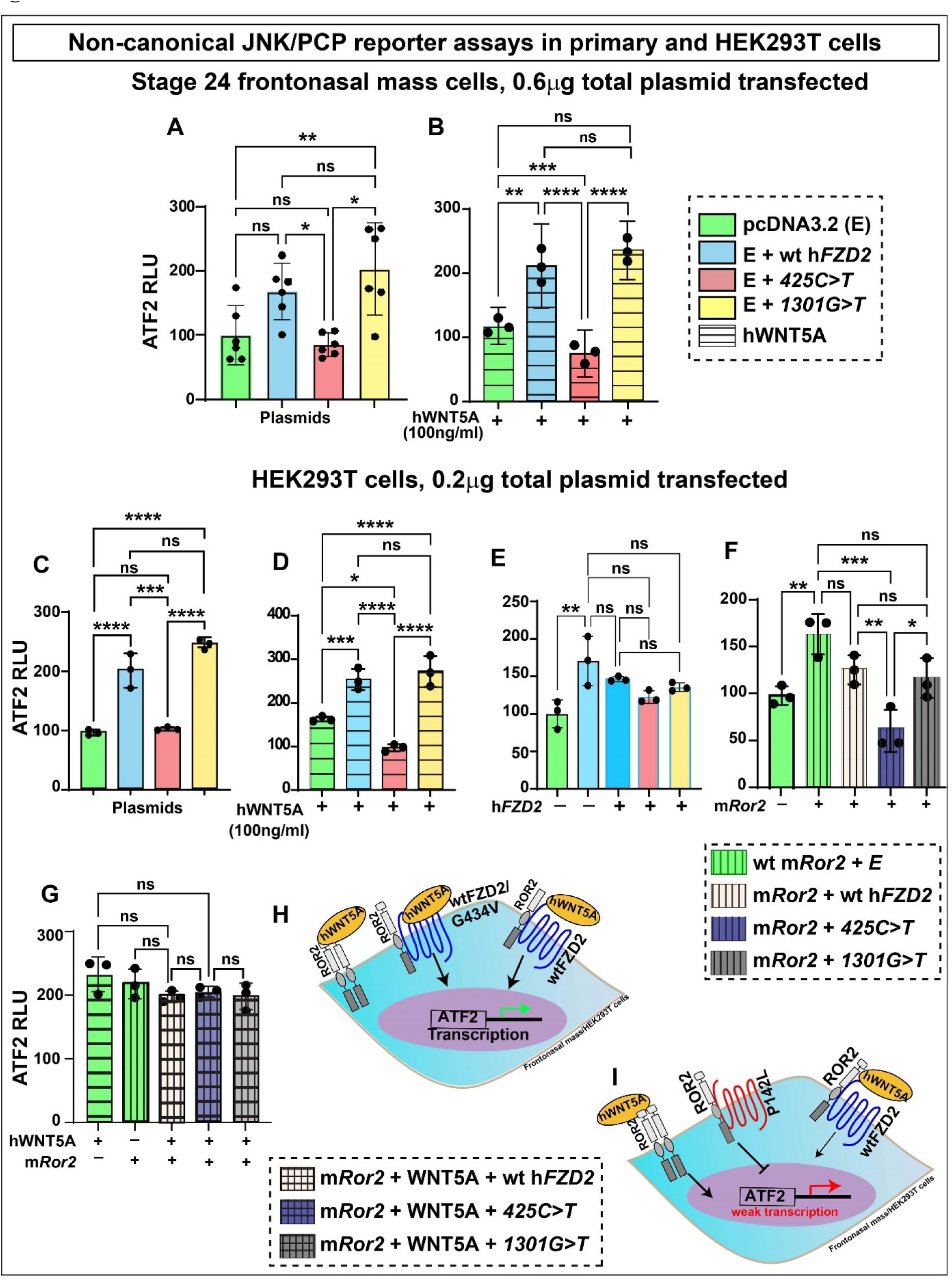
Effects of h*FZD2* variants of WNT non-canonical JNK-PCP pathway in frontonasal mass cultures and HEK293T cells. Activating transcription factor 2 (ATF2) used to measure noncanonical WNT JNK PCP pathway activity in stage 24 frontonasal mass micromass cultures or HEK293T cells. (A,C) The *425C>T* variant showed significantly lower ATF2 activity compared to wt h*FZD2* and *1301G>T*. (B,D) Recombinant human WNT5A (100ng/ml) was added 24 h post-transfection. wt h*FZD2* and *1301G>T* variant activated ATF2. The *425C>T* variant was significantly weak in activating ATF2 compared to wt h*FZD2* and *1301G>T*. (E) In HEK293T cells, all h*FZD2* variants combined in equimolar amounts with wt h*FZD2* (0.1ug wt + 0.1ug empty/wt/variant) showed comparable ATF2 activation. (F) When combined with m*Ror2*, the wt h*FZD2* and *1301G>T* plasmids showed similar ATF2 activation but *425C>T* significantly failed to activate ATF2. (G) Combination of h*FZD2*, m*Ror2*, and WNT5A showed comparable ATF2 activity for all h*FZD2*. (H) Schematic showing JNK-PCP pathway activity of wt h*FZD2* and *1301G>T* (I) and the *425C>T* variant. Three biological replicates, three technical replicates and the experiment was repeated three times. Statistics done with one-way ANOVA followed by Tukey’s post hoc test multiple comparison test using Prism 10.1.0. The error bars represent one standard deviation. Key: For A and B, 0.6µg of plasmid (0.3 µg of 0.3 µg of pcDNA3.2 + 0.3µg of plasmid *FZD2*) was used for transfection of frontonasal mass cultures; For C-F, 0.2µg of plasmid (0.1 µg of pcDNA3.2/ h*FZD2* + 0.1µg of m*Ror2*) was used for transfection of HEK293T cells; Key: RLU – relative luciferase activity, *≤0.05, **≤0.01, ***≤0.001, ****≤0.0001, ns – not significant.

There was no evidence of synergistic or antagonistic effects between the wtFZD2 and the variants in the JNK/PCP pathway unlike the STF results (Fig. 6E; 7E). FZD2 also binds to ROR2 to form heterodimers that transduce signals from WNT5A ligand (Cadigan and Waterman, 2012; Mikels et al., 2009; Nishita et al., 2010; Oishi et al., 2003). When plasmid containing mouse *Ror2* (m*Ror2*) was transfected into HEK293T cells, the ATF2 reporter was significantly activated compared to the empty vector (Fig. 7F). The ATF2 reporter activity in the presence of m*Ror2* was equivalent to wt*FZD2* and *1301G>T* (Fig. 7F). In contrast, in presence of *425C>T* and m*Ror2* plasmids, the ATF2 reporter failed to activate (no significant difference compared to pcDNA; Fig. 7F). While the Ror2-Fz7-Wnt5a complex has been shown experimentally to activate the JNK-PCP pathway (Nishita et al., 2010), the involvement of FZD2 in a complex with ROR2 and the WNT5A ligand has not been investigated until now. Both WNT5A and *Ror2* (Fig. 7G) on their own can effectively activate ATF2 (Fig. 7G)(Gignac et al., 2019; Gignac et al., 2023). When we combined hWNT5A and m*Ror2* with the wt or variant h*FZD2* variants, there was no further increase in ATF2 activity compared to WNT5A or m*Ror2* alone (Fig. 7G). This data aligns with that of others where homodimers of Ror2 were shown to form in the presence of WNT5A (Liu et al., 2008). Our data showed that when combined with WNT5A and *Ror2*, all three *FZD2* (wt or variant) showed no further increase in activity. This suggests that *FZD2* may have a minimal role in the WNT5A-ROR2-FZD2 complex and in activating the JNK/PCP pathway. This data is similar to previously published study which showed that binding of WNT5A to ROR2, prompts homodimerization of ROR2, excluding FZD from the signalosome (Feike et al., 2010; Liu et al., 2007; Liu et al., 2008).

In summary, our luciferase assays showed that that the *1301G>T* variant is weaker in activating the canonical pathway (Fig. 8B,C) but behaved like the wt *FZD2* in the ATF2-reporter assay (Fig. 7H, Fig. 8B,E). On the other hand, the *425C>T* variant is weak in activating the canonical (Fig. 8B,C) and shows a loss-of-function in the non-canonical JNK/PCP pathway (Fig. 7I, Fig. 8B,D). Both ADRS-*FZD2* variants have dominant-negative effects on the activity of wt h*FZD2* in the canonical β-catenin mediated pathway (Fig. 6H; 8D,E).

**Figure 8:**
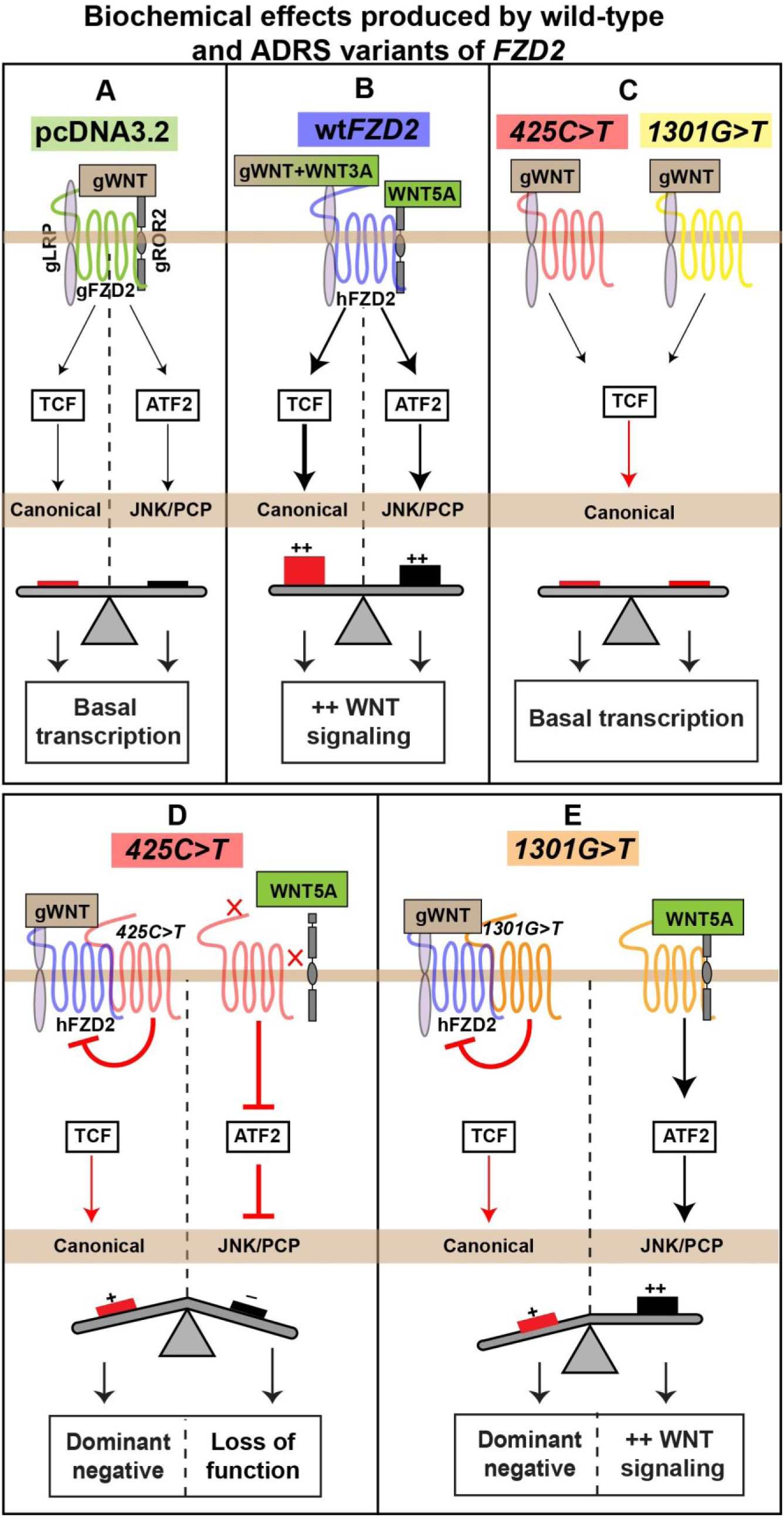
Summary of signaling effects produced by h*FZD2* or h*FZD2* variants. Schematic of biochemical activity of wt or variants of h*FZD2*. (A) Transfection of the empty vector (pcDNA3.2) in frontonasal mass mesenchyme or HEK293T cells showed basal level activity in both the canonical (STF, red) and JNK-PCP pathways (ATF2, black). (B) wt h*FZD2* (blue) significantly activated STF and ATF2. (C) The *425C>T* (red) and the *1301G>T* (yellow) variant weakly activate STF. (D) The *425C>T* variant when combined with wt h*FZD2* showed a dominant-negative on the activity of the wt. When *425C>T* was combined with WNT5A or m*Ror2*, JNK-PCP pathway was not activated. (E) The *1301G>T* variant also when combined with wt h*FZD2* showed dominant-negative effect. The *1301G>T* variant by itself and combined with WNT5A and m*Ror2* activated JNK-PCP pathway to the same extent as the wt h*FZD2*.

## DISCUSSION

Most of the genes that cause RS lie in the non-canonical WNT pathway except for *DVL* and *FZD2,* which also function in the canonical WNT pathway. There are ten *FZD* receptors in humans but interestingly only *FZD2* has been associated with pathogenesis of ADRS (Lima et al., 2022; White et al., 2018). In this study, we characterized two missense *FZD2* variants (*425C>T* and *1301G>T*) (White et al., 2018). We showed evidence of pathogenicity in cell signalling caused by both variants of *FZD2*. In addition, we discovered that embryonic facial morphogenesis was altered by the variants and these changes correlate with the broad nasal bridge and wide-set eyes of RS cases. We have clarified the causal relationship between *FZD2* variants (*425C>T* and *1301G>T*) and the craniofacial features of RS.

### Effects of *FZD2* variants on skeletogenesis

Robinow syndrome primarily affects the skeleton of the face, limbs, and vertebrae (Kaissi et al., 2020; Shayota et al., 2020). In this study, misexpression of wt or variant h*FZD2* in embryonic face did not cause morphological defects in the upper beak but inhibited patterning and ossification. These decreased ossification phenotypes are comparable to the effect of misexpression of ADRS h*WNT5A^248G>C^* variant that caused short and wide limbs and delay in ossification (Gignac et al., 2019). Likewise, in mouse model, an inducible transgene containing *Wnt5a* inhibited cranial bone formation (van Amerongen et al., 2012). On further investigation, we found that exogenous h*FZD2* downregulated WNT pathway mediator, β-catenin, at early embryonic stages. β*-catenin* is crucial for cell fate determination and osteoprogenitor specification during intramembranous bone formation (Goodnough et al., 2012). Loss of β*-catenin* in osteoblasts (*Collagen-Cre;*β*-catenin^floxed/floxed^*) leads to low bone-mass in mice (Glass et al., 2005). Additionally, conditional deletion of β*-catenin* in early osteogenic progenitor cells has been shown to prevent osteoblast differentiation (Day et al., 2005; Hill et al., 2005). Thus, our data showing that exogenous h*FZD2* caused downregulation of *CTNNB1* at early stages of development may lead to the hypoplasia of intramembranous bones.

Even thought in vivo data showed no differences in chondrogenesis between wt and variant h*FZD2*, in vitro micromass cultures revealed that the h*FZD2* variants have inhibitory effects on cartilage compared to wt at 8 days. The two variants were weaker in activating the canonical WNT pathway compared to wt in luciferase assays carried out at 3 days of culture. In contrast, at day 8 of culture, the variants caused ectopic activation of canonical WNT pathway. These temporal changes in levels of canonical WNT activity are not surprising. The later suppression of chondrogenesis is likely correlated with higher canonical WNT activity (Hill et al., 2005; Hosseini-Farahabadi et al., 2013). Furthermore, a second factor leading to inhibition of chondrogenesis in culture is the upregulation of a chondrogenic repressor, TWIST1 (Goodnough et al., 2012).

### The new role of FZD2 in controlling midfacial narrowing

Previous work from our lab showed that frontonasal mass narrowing at stage 28 is dependent on small-GTPase signaling mediated by ROCK (Danescu et al., 2021). In that study, we excluded oriented cell division or pressure from the expanding eyes as mechanisms for frontonasal mass narrowing. Indeed, the direct inhibition of ROCK with Y27632 prevents chicken embryo facial narrowing (Danescu et al., 2021). Our present data extends these results and finds two *FZD2* variants also modulate facial narrowing at the same stage of development. The *425C>T* variant has significantly weaker ATF2 reporter activity via the JNK-PCP pathway (JNK is downstream of RAC)(Qin et al., 2024). The ATF2 reporter was specifically designed to read out JNK-dependent activation of transcription factor ATF2 (Ohkawara and Niehrs, 2011). However, the *1301G>T* variant also inhibited facial narrowing without affecting the ATF2 reporter. The reason could be that the *1301G>T* variant inhibits the second PCP pathway – RhoA-ROCK(Qin et al., 2024). It is necessary in future to connect the h*FZD2 1301G>T* variant directly to measures of ROCK activity. All forms of ADRS have the same distinct facial features, hypertelorism and a broad nasal bridge (Zhang et al., 2022). Our data suggests that lower PCP activity in the frontonasal mass at a specific time in development may be the common mechanism.

### Is the BMP pathway involved in mediating RS phenotypes?

The crosstalk between the WNT and BMP pathways is crucial for embryonic bone development and postnatal bone homeostasis (Itasaki and Hoppler, 2010; Lin and Hankenson, 2011). In our study along with WNT pathway, we showed that the h*FZD2* variants caused a parallel downregulation of *BMP2*, *BMP7*, the BMP antagonist *NOGGIN*. Consistent with this data on BMP expression reduction, we found that *425C>T* variant failed to activate the SOX9-luc reporter. An interesting finding was downregulation of BMP regulator *BMPER* (a homologue of *crossveinless* in Drosophila)(Conley et al., 2000), was consistently downregulated by all h*FZD2*. *Bmper^-/-^* mice have severe skeletal defects (Ikeya et al., 2006) and downregulation of *BMPER* has been shown to inhibit osteogenic differentiation in human bone marrow stem cells (Xiao et al., 2018). Furthermore, loss-of-function variants in *BMPER* cause a rare congenital skeletal disorder – Diaphanospondylodysostosis (Arredondo Montero et al., 2022; Braun et al., 2021; Greenbaum et al., 2019). Overall, the h*FZD2* variants had severe effects on the BMP pathway compared to wt. This is the first study to establish a link between ADRS-*FZD2* variants and the BMP pathway. Future studies should consider participation of the BMP pathway in mediating RS phenotypes.

### Micromass cultures clarify variant-specific effects in signaling and differentiation

In our study there are certain experiments that do not show selective effects of the variant h*FZD2* as compared to wt*FZD2* and these support a general gain of function due to presence of exogenous *FZD2*. This includes the in vivo ossification phenotypes, gene expression changes observed in qRT-PCR, BrdU and TUNEL assays. We also have presented data that show selective effects of the variants in vivo (increased frontonasal mass width), luciferase assays (STF, ATF, SOX9) and micromass cultures (reduced thickness and amount of cartilage, increased TWIST1 and CTNNB1 expression). Taken together, the strongest evidence for changes in variant function are those that were performed on primary facial mesenchyme cells in micromass cultures and in luciferase assays. Excluding the epithelium was helpful in identifying the effects of the variants on facial mesenchyme that could not be appreciated in vivo.

### Over-expression in chicken embryos complements mouse models for RS variants

The over-expression experiments in chicken embryos tests whether a variant is sufficient to change the course of development. The results are complementary to mouse genetic models where the variant is knocked into the equivalent location of the genome. In the mouse constitutive knock-in model, the spatio-temporal regulation of gene expression is conserved and mirrors the levels of expression seen in the human AD heterozygous cells. Indeed, the Fzd2 knock-in model was recently published (Liegel et al., 2023). In this study, the *Fzd2^W548*^* variant caused 100% lethality in the embryos, even in the heterozygous state. It was necessary to go to great lengths to produce F0 embryos including electroporation of the oviduct. Since animals could not be bred, there were limits to the number of mutant embryos available to study at different stages of development. Therefore, even though in theory the mouse knock-in model seems superior unexpected problems can occur. There are other examples of lethality in heterozygous mice with constitutively expressed mutation such as this one with a missense mutation knocked into *Ifitm5* which in humans causes type V osteogenesis imperfecta (Rauch et al., 2018). The mouse models may also have no phenotype unless bred to homozygosity. Therefore, with all these caveats about the mouse model, overexpression in the chicken embryo or other animal models (Gignac et al., 2023) is required to fully define the effects of the mutation. Furthermore, our chicken data suggests that other variants in *FZD2* may be worth following up in the mouse based on their signaling defects and effects on cell differentiation.

Overall, this study provides mechanistic understanding of the developmental, molecular, cellular, and biochemical processes that are affected in patients carrying mutations in *FZD2*. These discoveries broaden our understanding of the WNT pathway and will be useful in future studies aimed at developing therapeutic interventions for ADRS patients.

## MATERIALS AND METHODS

### Chicken embryo model

White leghorn eggs (*Gallus Gallus*) obtained from the University of Alberta were incubated to the appropriate embryonic stages, based on the Hamilton Hamburger staging guide (Hamburger and Hamilton, 1992). All experiments were performed on prehatching chicken embryos which are exempted from ethical approval by the University of British Columbia Animal Care Committee and the Canadian Council on Animal Care.

### Cloning of human *FZD2* constructs

The open reading frame encoding human *FZD2* sequence was purchased from GeneCopoeia (Rockville, MD, clone #GC-S0193-B). Restriction-free cloning (Bond and Naus, 2012) was used to add a C-terminal Flag-tag (DYKDDDDK) into the wt h*FZD2* vector (pShuttle) and a stop codon was added at the 3’ end of the tag. Two autosomal dominant Robinow syndrome associated missense variants (*1301G>T*, *425C>T*) were knocked into the Flag-tagged wt h*FZD2* (pShuttle) vector (Table S6). Flag-tagged wt or variant *FZD2* vectors (pShuttle) were then recombined into destination vectors (pcDNA 3.2 expression vector and RCAS retroviral vector) using Gateway LR clonase II enzyme mix (Thermo Fisher #11791019)] as described (Hosseini-Farahabadi et al., 2017). *GFP* was not fused to the 5’ end of FZD2 constructs due to the large size of *GFP* interfering with localization of FZD2 to the cell membrane. The small size of the FLAG-tag and its high hydrophilicity, tend to decrease the possibility of interference with protein expression and function (Einhauer and Jungbauer, 2001). To create the RCAS viruses, a Gateway compatible RCASBPY destination vector was used for recombination with the pShuttle vector using LR Clonase II(Loftus et al., 2001). It should be noted that the insert size for RCAS is restricted to 2.4 kb (Loftus et al., 2001), therefore a *GFP*-tag (730 bp) to track the viral spread could not be added. Viral constructs are available on request to the senior author.

### Growth of RCAS viral particles and viral titre

Replication-Competent ASLV long terminal repeat (LTR) with a Splice acceptor (RCAS) (Hughes, 2004; Loftus et al., 2001) (Hughes, 2004)plasmid DNAs (2.5µg) encoding *GFP*, human *FZD2*, or two *FZD2* variants (425C>T and 1301G>T) were transfected into the DF-1 immortalized chicken fibroblast cell line (American Type Culture Collection, CRL-12203) using Lipofectamine 3000 (Thermo Fisher #L3000-008) following the manufacturer’s guidelines. RCAS virus containing *GFP* insert (kindly provided by A. Gaunt) served as a control in virus overexpression studies, as published(Geetha-Loganathan et al., 2014; Gignac et al., 2019; Gignac et al., 2023; Hosseini-Farahabadi et al., 2013; Hosseini-Farahabadi et al., 2017). DF1 cells were cultured at 37°C and 5% CO_2_ in DMEM (ThermoFisher#1967497) medium supplemented with 10% fetal bovine serum (FBS; Sigma #F1051) and 1% penicillin/streptomycin (ThermoFisher #15070-063). The cells were maintained in 100 mm culture dishes with media changes every other day and passaged 1:2 two to three times per week using trypsin-EDTA (0.25%, ThermoFisher #25200-072). After six weeks of culturing, the viral particles were collected and centrifuged in a swing bucket SW28 rotor (Beckman #97U 9661 ultracentrifuge) for 2.5 hours (no brake) at 25,000 rpm at 4°C. The supernatant was carefully removed, and the resulting pellet was resuspended in 50-100 µl Opti-MEM (ThermoFisher#319850962). This suspension was then incubated overnight at 4°C. The concentrated viral particles obtained were aliquoted (5µL aliquots), rapidly frozen in methanol + dry ice, and stored at −80°C for future use (Goodnough et al., 2012).

To determine viral titer, 50-60% confluent DF1 fibroblasts were infected with serial dilutions of 2μl of concentrated viral stock. After 36 hours of virus incubation, cells were fixed in 4% paraformaldehyde for 30 mins. Immunocytochemistry with Group Associated Antigens (GAG) antibody was performed on virus-treated cells (Table S9). The cells were permeabilized for 30 mins with 0.1% Triton X-100, followed by blocking in 10% goat serum and 0.1% Triton X-100 and overnight incubation with the primary antibody (Table S9). Fluorescence images were captured using a Leica inverted microscope at 10x with a DFC7000 camera. The analysis of viral titer was done with ImageJ’s cell counter tool by determining the proportion of GAG-positive cells per mL in a 35mm culture plate. Virus titer = # of GAG positive cells * (Area counted * total cells expressing GAG)/2*1000 (Fig. S7).

### Chicken embryo injections

Fertilized eggs obtained from the University of Alberta, Edmonton, were incubated in a humified incubator at 38°C until Hamilton Hamburger (Hamburger and Hamilton, 1951; Hamburger and Hamilton, 1992) stage 15 (E2.5). Concentrated RCAS (titer = >2 x 10^8^ IU/mL) retrovirus viral particles (∼5μl) combined with Fast Green FCF stain (0.42%, Sigma #F7252) (1μl) were injected into the frontonasal mass (anatomic region bounded by the nasal slits) of stage 14-15 chicken embryos (25-28 somites) using glass filament needles (thin-wall borosilicate capillary glass with microfilament, A-M systems #615000) and a Picospritzer microinjector (General valve corp. #42311). The infection of embryos with RCAS at stage 15 (E2.5) was performed to ensure maximum infection of facial prominences(Geetha-Loganathan et al., 2009; Hosseini-Farahabadi et al., 2017). Due to accessibility, all injections were made into the right frontonasal mass as the chick embryos turn on their left side during development. The facial prominences form around stage 20, and the complex and temporally regulated patterning occurs between stages 20-29. The skeletal derivatives of the frontonasal mass are fully patterned and ossified between stages 34-40. The investigation encompassed multiple embryonic stages to comprehensively analyze these developmental processes. After conducting the overexpression of high-titer *FZD2* viruses at stage 15, the retrospective determination of virus location was carried out on histological sections. These sections were stained with Group-associated antigens (GAG), an antibody that identifies specific proteins in the RCAS virus (Geetha-Loganathan et al., 2009; Gignac et al., 2023; Hosseini-Farahabadi et al., 2017) or an antibody against the FLAG sequence (Gignac et al., 2023; Tetenborg et al., 2020) (Table S9).

### Wholemount staining of skulls

To study skeletal elements, embryos were grown until stage 38 (10 days post-injection, Table S3). The embryos were washed in 1x phosphate-buffered saline (PBS; 137 mM NaCl, 8.1 mM Na_2_HPO_4_, 2.7 mM KCl, 1.5 mM KH_2_PO_4_; pH 7.3) and fixed in 100% ethanol for 4 days. After removal of eyes and skin, the embryos were transferred to 100% acetone for another 4 days. Subsequently, the heads were stained with a freshly prepared bone and cartilage stain (0.3% Alcian blue 8GX (Sigma #A5268) in 70% ethanol, and 0.1% alizarin red S (Sigma # A5533) in 95% ethanol, with 1 volume of 100% acetic acid and 17 volumes of 70% ethanol) for two weeks on shaker at room temperature. Following staining, the skulls were washed in water and cleared in a 2% KOH/20% glycerol solution on shaker for 4 days, followed by immersion in 50% glycerol for imaging. The heads were stored in 100% glycerol post-imaging. Phenotyping was conducted by photographing the right lateral, superior and palatal views of cleared heads using a Leica DFC7000T microscope camera. Skeletal preparations from each virus type were analyzed for changes in the size or shape of bones derived from the frontonasal mass, missing bones, or qualitative reduction in ossification observed as reduced alizarin red stain. Statistical analysis was performed using contingency analysis Chi-square test in GraphPad Prism 10.1.0.

### Primary cultures of frontonasal mass mesenchyme

Stage 24 chicken embryos were extracted from the eggs and their extra-embryonic tissues were removed in cold phosphate-buffered saline (PBS). The frontonasal mass was dissected in cold Hank’s balanced saline solution (HBSS) (without calcium and magnesium) (ThermoFisher #14185052) with 10% FBS and 1% Antibiotic-Antimycotic (Life Technologies #15240-062). Dissected frontonasal pieces were incubated in 2% trypsin (Gibco) at 4°C for 1 hour. Hank’s solution was added to inhibit the enzymatic activity of trypsin. Ectoderm was manually peeled off from the frontonasal mass pieces. The cell solution was then centrifuged at 1000 g, 4°C for 5 minutes. The supernatant was removed and the frontonasal mass pieces were resuspended in Hank’s solution. The mesenchymal cells were counted using a hemocytometer and 2×107 cells/ml were resuspended in chondrogenic media (micromass media) containing DMEM/F12 medium (Corning #10-092-CV) supplemented with 10% FBS, 1% L-Glutamine (ThermoFisher #25030), Ascorbic acid (50 mg/ml) (ThermoFisher #850-3080IM), 10 mM β-glycerol phosphate (Sigma Aldrich #G9422) and 1%, Antibiotic-antimycotic (ThermoFisher #15240-062). The cells in suspension were subsequently infected with 3µL *GFP* (control), wt h*FZD2*, or h*FZD2* (*1301G>T* or *425C>T*) variant containing viruses. The 10µl of cells suspension infected with virus was plate micromass cultures (3-4 spots per 35mm culture dish, NUNC #150318) at a density of 2 x 10^7^ cells/ml, (Hosseini-Farahabadi et al., 2013; Richman and Tickle, 1989; Richman and Tickle, 1992; Underhill et al., 2014). The culture plates were incubated at 37 °C and 5% CO_2_ for 90 minutes to allow cells to attach and then flooded with 2 ml of micromass media. Thereafter, micromass culture media was changed every other day for experimental time points of day 4, 6, and 8.

### Wholemount staining of micromass cultures

On day 4, 6, and 8, cultures were fixed in 4% paraformaldehyde for 30 minutes at room temperature and subjected to wholemount staining. To detect the mineralization of cartilage using alkaline phosphatase stain (Table S8), fixed cultures were incubated at room temperature in 100mM Tris for 30 minutes (pH 8.3). Following this, the cultures were stained with 0.5% Alcian Blue in 95% EtOH: 0.1 M HCl (1:4) to detect the area occupied by cartilage, as previously described (Hosseini-Farahabadi, 2013; Underhill, 2014). All cultures were counterstained with 50% Shandon’s Instant Hematoxylin (ThermoFisher #6765015). The stained cultures were photographed under standard illumination using a stereomicroscope (Leica #M125). Wholemount staining was conducted on three biological and three technical replicates, and the experiment was repeated five times.

### Histology and Immunofluorescence

Embryos collected at stage 28 (Table S5) or micromass cultures (day 4, 6, 8, Table S7) were fixed in 4% PFA. The embryo samples were immersed in the fixative for 2-3 days at 4°C. The RCAS-infected cultures fixed in 4% paraformaldehyde for 30 mins. The cultures were then removed from the plate using a cell scraper (ThermoFisher #08-100-241), embedded in 2% agarose (Sigma #A9539) on a cold ice slab, and subsequently wax-embedded. The embryos (positioned frontally) and cultures (positioned transversely) were embedded in paraffin wax and sliced into 7µm sections. The sections were then utilized for histological and immunostaining analysis.

Selected frontal (embryos) and transverse (micromass cultures) sections were stained to visualize the differentiated cartilage and bone. Sections were dewaxed in xylene, rehydrated from 100% ethanol to water, and stained with 1% Alcian blue 8GX (in 1% acetic acid) for 30 minutes. After staining, sections were rinsed in 1% acetic acid and water. Subsequently, sections were stained in Picrosirius Red (0.1% Sirius Red F3B in saturated picric acid) for 1 hour in dark, followed by rinsing in 1% acetic acid, dehydration through ethanol, back to xylene, and mounted with Shandon Consul-mount (Thermo Scientific #9990441).

Immunofluorescence analysis was conducted on in vivo and day 6 and 8 cultures. Specific antibodies and treatments performed for each assay are outlined in Table S9. Primary antibodies were allowed to incubate overnight at 4°C, while secondary antibodies were incubated at room temperature for 1.5 hours unless otherwise specified. Sections were counterstained with Hoechst (10μg/ml #33568, Sigma) and incubated for 30 minutes at room temperature, then mounted with Prolong Gold antifade (ThermoFisher #P36930). Fluorescence images were captured using 20X objective on a slide scanner (3DHISTECH Ltd., Budapest, Hungary).

### Apoptosis and Cell proliferation

Apoptosis was analyzed using TUNEL (Terminal deoxynucleotidyl transferase dUTP nick end labeling) assay on sections obtained from virus infected frontonasal mass stage28 and micromass cultures sections day 6 and day 8. The TUNEL assay was performed using ApopTag Plus *in Situ* Apoptosis Fluorescein Detection Kit (Millipore Sigma # S7111).

For cell proliferation studies, embryos at stage 28, stage 29 or stage 30 were labeled with 50µl of 10 mM BrdU (Bromodeoxyuridine; Sigma #B5002) and incubated at 38°C for 1 hour prior to euthanizing. For labeling micromass cultures, 50µl of 10mM BrdU was added to the culture media (37°C 5% CO_2_) 1 hour before fixing day 6 and day 8 cultures in 4% PFA. Immunostaining was performed on the sections with anti-BrdU (Developmental Studies Hybridoma bank, 1:20, #G3G4) as described in Table S9. Fluorescence images were collected with a 20X objective on a slide scanner (3DHISTECH Ltd., Budapest, Hungary).

### qRT-PCR on frontonasal mass in vivo and in vitro

Viral spread in the frontonasal mass was quantified using primers specific to human *FZD2* (primer set, Table S10). Three biological replicates containing five to six pieces of the right half of the frontonasal mass pooled in each sample were harvested for each virus at stage 28 (E5.5) and stage 30 (E7.5). Similarly, three biological replicates containing pools of twelve micromass cultures per replicate were collected on day 6 and day 8. Total RNA was isolated from frontonasal masses using Qiagen RNAeasy kit (#75144, Toronto, Canada). Sybr green-based quantitive reverse transcriptase polymerase change reaction (Advanced universal SYBR® Green supermix; Bio-Rad #1725271) (qRT-PCR) was carried out using an Applied Biosystems StepOnePlus instrument. qRT-PCR cycling conditions were - 95°C for 10min, 40X (95°C for 5s, 60°C for 20 seconds). Analysis used human-specific primers for *FZD2* and avian primers were used (Table S10). The expression of each biological replicate was normalized to 18s RNA (Applied Biosystems, 4328839) and then these ΔCt values were used to calculate the ΔΔCt relative to the average levels of expression of the gene in *GFP*-infected cultures. The ΔΔCt method was used to calculate relative fold-change expression between RS-*FZD2* infected frontonasal mass and *GFP* as described (Schmittgen and Livak, 2008). Statistical analysis was done with one-way ANOVA Tukey’s post hoc test in GraphPad Prism 10.0.2. A sample size calculator was used to determine how many samples would need to be included in order to detect a P value of 0.05 80% of the time and it was necessary to collect 13 biological replicates. This number of biological replicates was not feasible for these studies.

### Luciferase reporter assays

Transient transfections for luciferase assays were performed in HEK293T (0.15 x 10^6^ cells/ml) (Gignac et al., 2019; Gignac et al., 2023) or untreated stage 24 frontonasal mass mesenchymal micromass cultures (1 x 10^7^ cells/ml) as described (Geetha-Loganathan et al., 2014; Hosseini-Farahabadi et al., 2013). Cells were transfected with Lipofectamine 3000 (Invitrogen, L3000-008, Nunc 24-well plates #142475) transfection reagent. HEK293T cells were transfected 24h after plating (40-50% confluence). Micromass cultures (2 x 10^7^ cells/ml) were allowed to attach for 45 mins after plating and transfection reagents were added to the culture spot 30 mins prior to flooding the culture plate with micromass media. The following plasmids were used: control/empty (pcDNA3.2/V5-DEST), h*FZD2*, FZD2*^425C>T^*, FZD2*^1301G>T^*, m*Ror2* (Addgene, #22613), single or in combinations (totalling to 0.2μg for HEK293 cells and 0.6μg for frontonasal mass cultures). Firefly reporter plasmids: SuperTopflash (STF; 0.2ug, Addgene plasmid #12456) and Activating Transcription Factor 2 (0.4ug; ATF2) (Ohkawara and Niehrs, 2011) along with Renilla luciferase was transfected for normalization (0.01μg). Recombinant Wnt3a (100 ng/ml, R&D, #5036-WN-010) or Wnt5a protein (100 ng/ml, R&D # 645-WN-010) was added 24h post transfection.

To measure the WNT canonical/β-catenin pathway and chondrogenic activity in micromass, the cultures were infected with 3μl of concentrated virus at the time of plating. 24h after plating, we performed transient transfection with either STF or SOX9 luciferase (pGL3 4×48) (Weston et al., 2002). Renilla luciferase was used as normalization control. Assay reading was done 48h after transfection representing day 3 of culture. The dual-luciferase reporter assay system (Promega #E1910) was used for all luciferase assays as described (Geetha-Loganathan et al., 2014). Luminescence activity was detected with a PerkinElmer Victor X2 Multilabel Microplate Reader at 1 s reading with OD1 filter. All data shown represents two to three independent experiments with three technical and three biological replicates carried out for each transfection mixture. Statistical analysis done using one-way ANOVA, Tukey’s post hoc test in GraphPad Prism 10.0.2. The number of biological replicates was determined by our previous studies using luciferase assays (Gignac et al., 2019; Gignac et al., 2023).

### Image analysis and statistics

1. For measurement of the width of the frontonasal mass at stage 28, distance between the nasal slits (illustrated in Fig. 2X) was measured manually with linear measurement annotation tool in CaseViewer (version 2.4). The data was analyzed using one-way ANOVA, Tukey’s test in GraphPad Prism 10.1.0.
2. The thickness of day 6 and 8 micromass cultures was measured using histological sections stained with Alcian blue and picrosirius red. The linear measurement annotation tool in CaseViewer (version 2.4) was utilized. For each experimental condition (day 6, day 8), measurements were performed on three cultures per virus type (biological replicates). Each biological replication represents an average of three ribbons. The data was analyzed using one-way ANOVA, Tukey’s test in GraphPad Prism 10.1.0.
3. For analysis of immunofluorescence staining performed on stage 28 samples (n=3), cells in the prenasal cartilage in all treated samples were counted in a 100 x 100µm^2^ area to get the average cell density. The right frontonasal mass was divided into four regions (100 x 250um^2^) (Fig. 2U) to count the proportion of cells expressing BrdU and TUNEL. All cell counts were performed twice with the counter plugin in ImageJ by a blind observer. The data was analyzed using one-way ANOVA, Tukey’s test in GraphPad Prism 10.1.0. Similar samples sizes were used in other studies on BrdU labelling (Gignac et al., 2019; Gignac et al., 2023).
4. For micromass cultures, the proportion of cells expressing TWIST1 and nuclear β-catenin, or cells labelled in the TUNEL assays, were counted manually by the first author and a blind observer within a 500 x 200µm² area. The total number of TWIST1/nuclear β-catenin, or TUNEL positive cells was divided by the total number of Hoechst+ cells to obtain the proportion of labelled cells. All cell counts were performed by the first author and by a blinded observer with the counter plugin in ImageJ. Statistical analysis was done with one-way ANOVA with Tukey’s post hoc test in GraphPad Prism 10.1.0.

## Supporting information

Tophkhane et al. Supplementary Data

## Acknowledgements

We are grateful to the Developmental Studies Hybridoma Bank (IA, USA) for providing antibodies. We thank Dr. Janel Kopp for use of the 3DHistech slide scanner for all chicken histology images. We acknowledge the excellent cell counting carried out by Dima Lim and Karen Chen.

## Author contributions

Conceptualization: S.S.T and J.M.R. Methodology: S.S.T and K.F.; Formal analysis: S.S.T. and J.M.R. Investigation: S.S.T., K.F., J.M.R.; Resources: J.M.R.; Writing - original draft: S.S.T. and J.M.R.; Writing - review & editing: S.S.T., J.M.R, E.M.V; Supervision: J.M.R.; Project administration: J.M.R.; Funding acquisition: E.M.V., J.M.R.

## Data availability

All relevant data can be found within the article and its supplementary information. The senior author will provide additional data and DNA constructs on request.

