## Supplementary material for "Craniofacial studies in chicken embryos confirm the pathogenicity of Frizzled2 variants associated with Robinow syndrome": Tophkhane et al. Supplementary Data

**Supplementary figures**

**Figure S1**

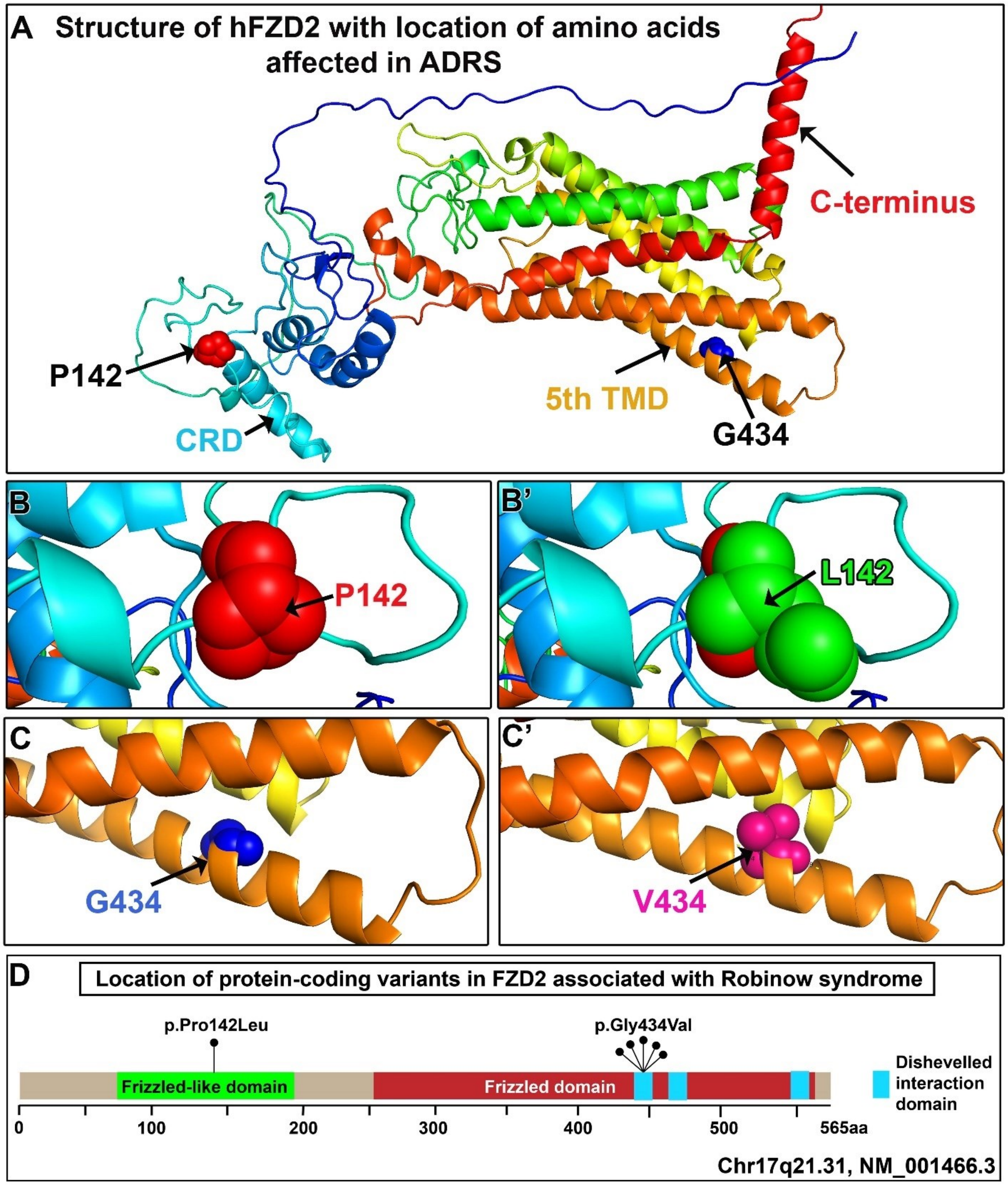

**Figure S1:Schematic representation of ADRS-associated FZD2$^{425C>T}$ and FZD2$^{1301G>T}$ variants**

FZD2$^{425C>T}$ codes for P142L and FZD2$^{1301G>T}$ codes for G434V. (A) The predicted ribbon structure of FZD2 protein created in PyMOL software using AlphaFold (Identifier# AF-Q14332-F1). The location of affected amino acids Proline 142 (P142 red spheres) in the cysteine rich domain (cyan) and Glycine 434 (G434 royal blue spheres) in the 5$^{th}$ transmembrane domain (TMD) (orange) are shown. The C-terminus is represented by red. (B') Magnified view of the Proline at position 142 (red spheres) (B) replaced by Leucine (green spheres). (C') Magnified view of Glycine at position 434 (blue spheres) (C) replaced by Valine (pink spheres). (D) Representation of the known functional domains of FZD2 (green, red) and the dishevelled interaction domain (turquoise) are illustrated. The predicted location of the ADRS missense mutations p.Pro142Leu (c.425C>T) and p.Gly434Val (c.1301G>T). The number of patients classified as ADRS patients are shown with black dots (modified from White et al., 2018; Zhang et al., 2022).

**Figure S2**

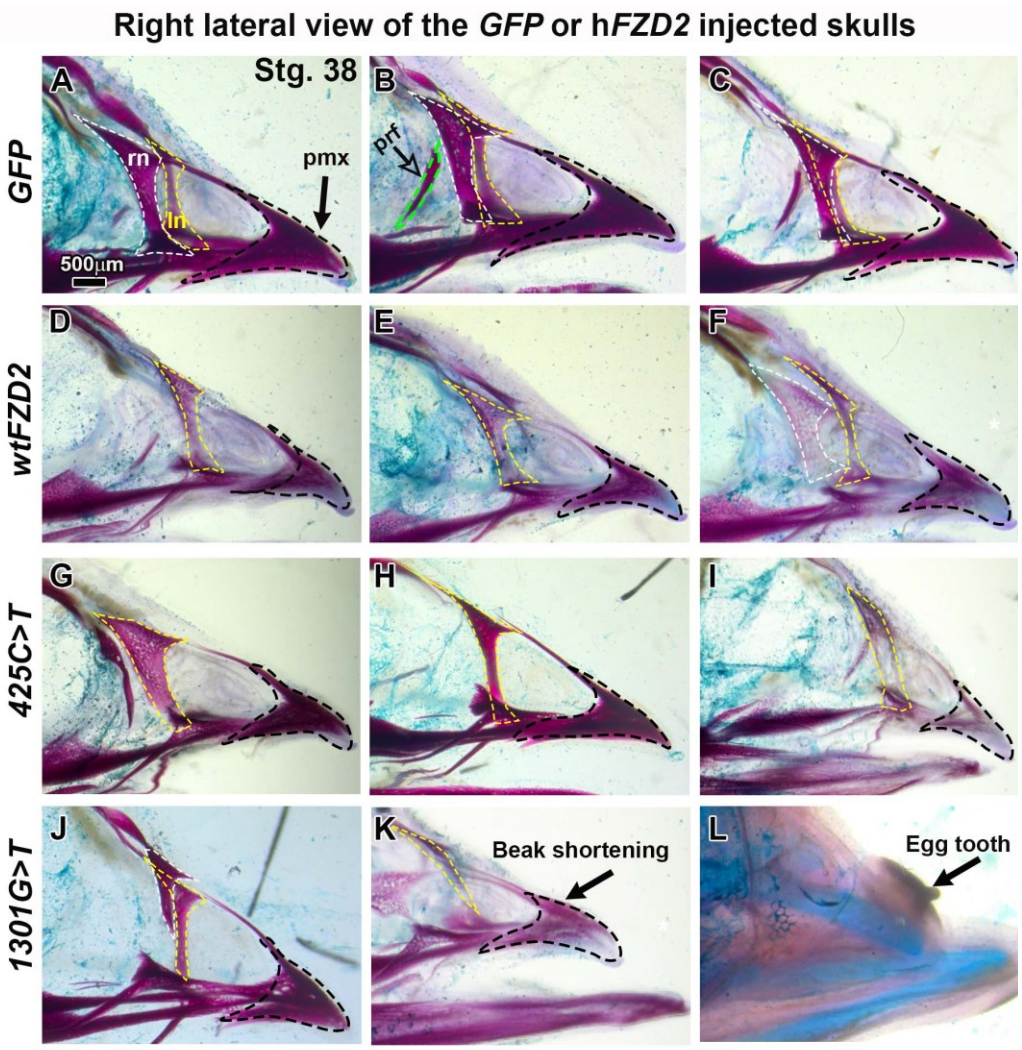


**Figure S2 Skeletal preparation of additional embryos injected with GFP, wild-type hFZD2 or mutant hFZD2 variants**

Embryos injected *GFP*, wild-type h*FZD2*, h*FZD2*$^{425C>T}$, or h*FZD2*$^{1301G>T}$ into the frontonasal mass at stage 15 (E2.5) and fixed 10 days post-injection at stage 38. (A-L) Wholemount skulls cleared and stained with alcian blue (cartilage) and alizarin red (bone). (A-C) *GFP* virus allowed normal patterning and ossification of frontonasal mass derived bones including premaxillary, nasal and prefrontal. (D-L) Embryos injected with wild-type h*FZD2* or mutant h*FZD2* viruses had no defects in the upper beak morphology but had missing nasal bone on the injected side (white dashed line – right nasal bone) while the nasal bone was present on the left side (yellow dashed line). The prefrontal bone (green dashed line) was absent in >90% of embryos and the premaxillary bone (white dashed line) was present but showed faint alizarin red staining. The size of the premaxilla appeared to be smaller compared to *GFP* controls. (K, L) Some embryos injected with h*FZD2*$^{1301G>T}$ had beak shortening and deviation (6/19). (L) An embryo with unossified skull and short upper beak injected with h*FZD2*$^{1301G>T}$. The faint cartilage stain (blue) in all skulls is a technical error. Key: ln, left nasal; pmx, premaxilla; prf, prefrontal; rn, right nasal. Scale bar: A-L= 2mm.

**Figure S3**

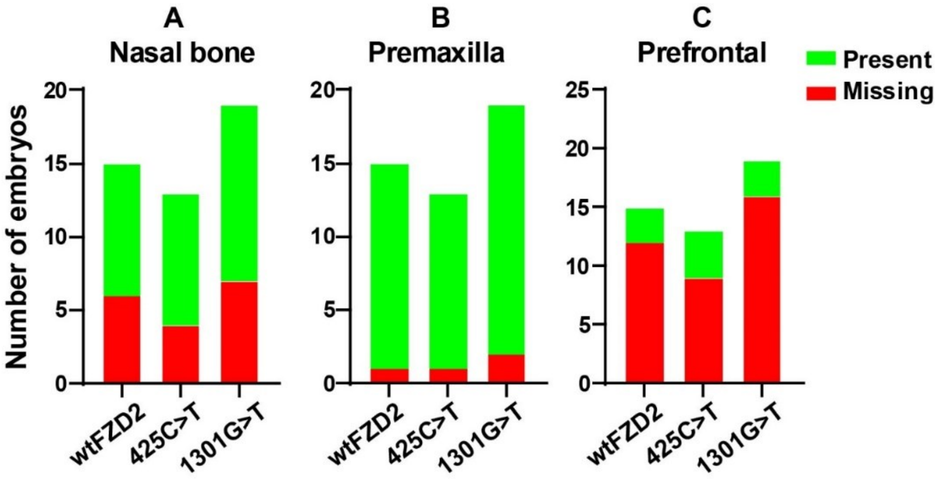


**Figure S3: Analysis of missing bones in embryos injected with h*FZD2* viruses**

Contingency analysis (Fisher's exact test) showing total embryos with normal or missing (A) nasal (B) Premaxillary, or (C) Prefrontal bones at stage 38 in vivo.

**Figure S4**

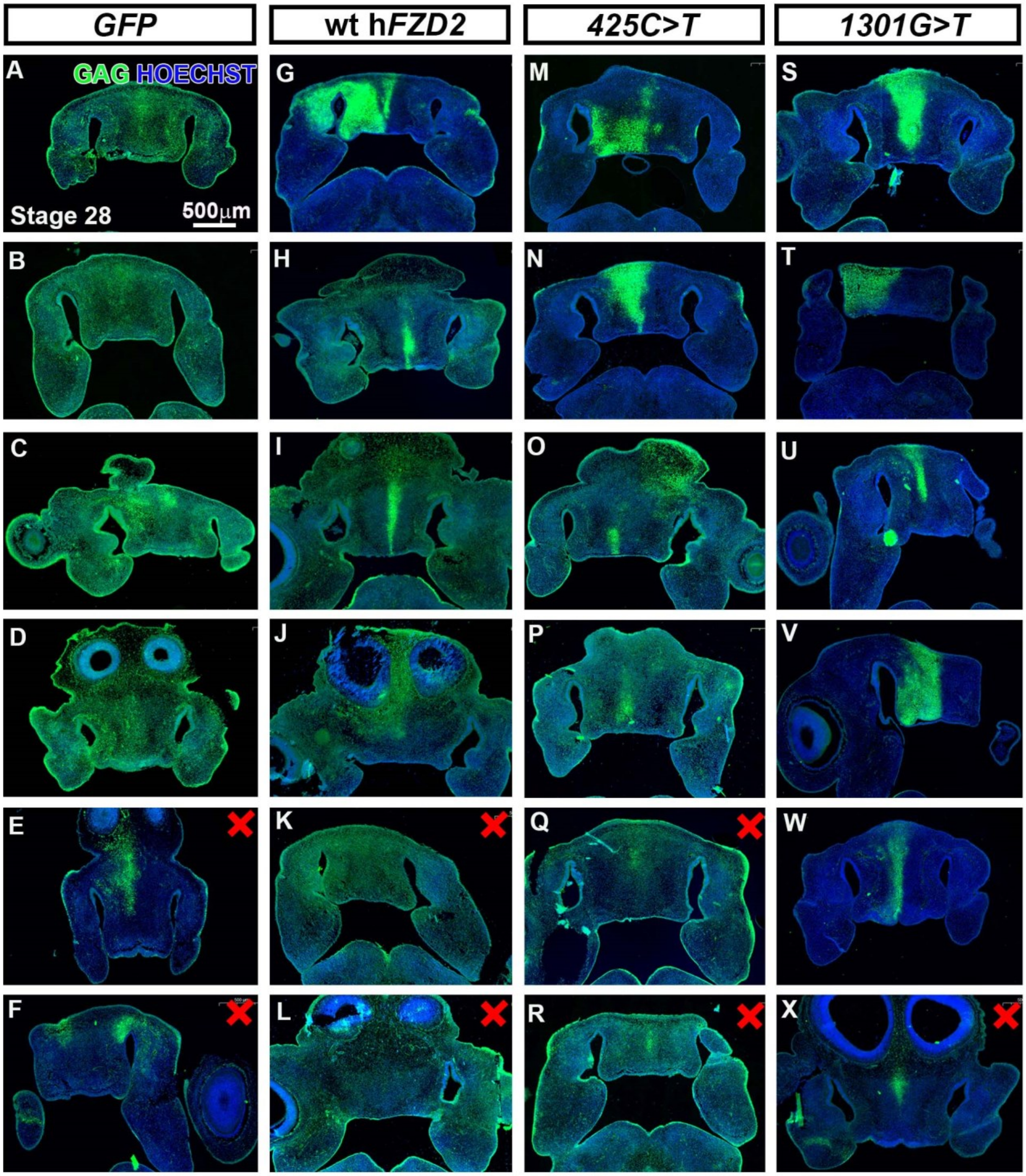


**Figure S4: Embryos at stage 28 (E5.5) injected with *GFP*, wt h*FZD2* or h*FZD2* variants**

Paraffin embedded serial sections were stained with anti-GAG antibody (green fluorescence) and counter stained with Hoechst (blue) to detect the viral spread in the frontonasal mass in embryos injected with (A-F) *GFP* (G-L) wt h*FZD2*, (M-R) *425C>T*, (S-X) *1301G>T* viruses. Embryos with lower GAG-stanning (red cross) were not used for molecular analysis. Scale bar = 500µm

**Figure S5**

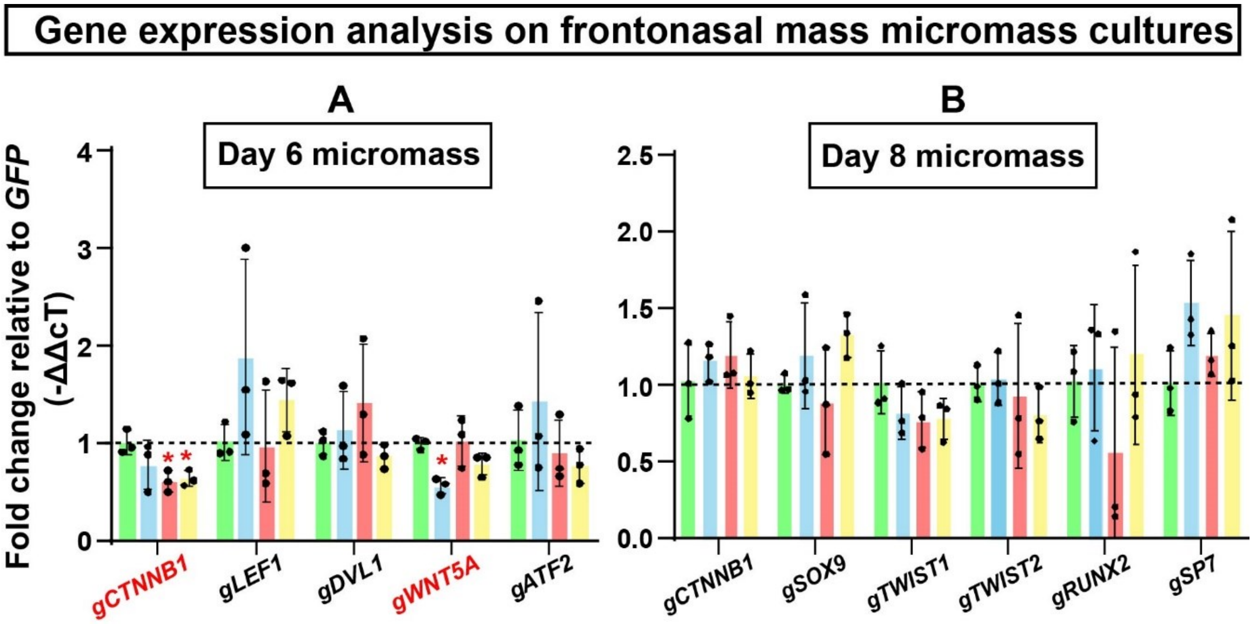


**Figure S5: qRT-PCR analysis showing effects of h*FZD2* viruses on mediators of the WNT pathway and skeletogenic mediators**

RNA-isolated from day 6 or day 8 micromass cultures, three biological replicates per virus type containing pool of 8-9 micromass cultures per biological replicate. The expression of each biological replicate was normalized to 18s RNA and then these ΔCt values were used to calculate the ΔΔCt relative to the average levels of expression of the gene in *GFP*. The graphs showing gene expression changes on (A) day 6 and (B) day 8. All statistical analysis was done using one-way ANOVA, Dunnett's multiple comparison test in GraphPad Prism 10.1.0. The error bars represent one standard deviation. Genes showing statistically significant differences are shown in red text. Black dashed line represents control *GFP*. Key: Each black dot represents a biological replicate. The p values are displayed with red asterisk on the graph. * - $p < 0.05$.

**Figure S6**

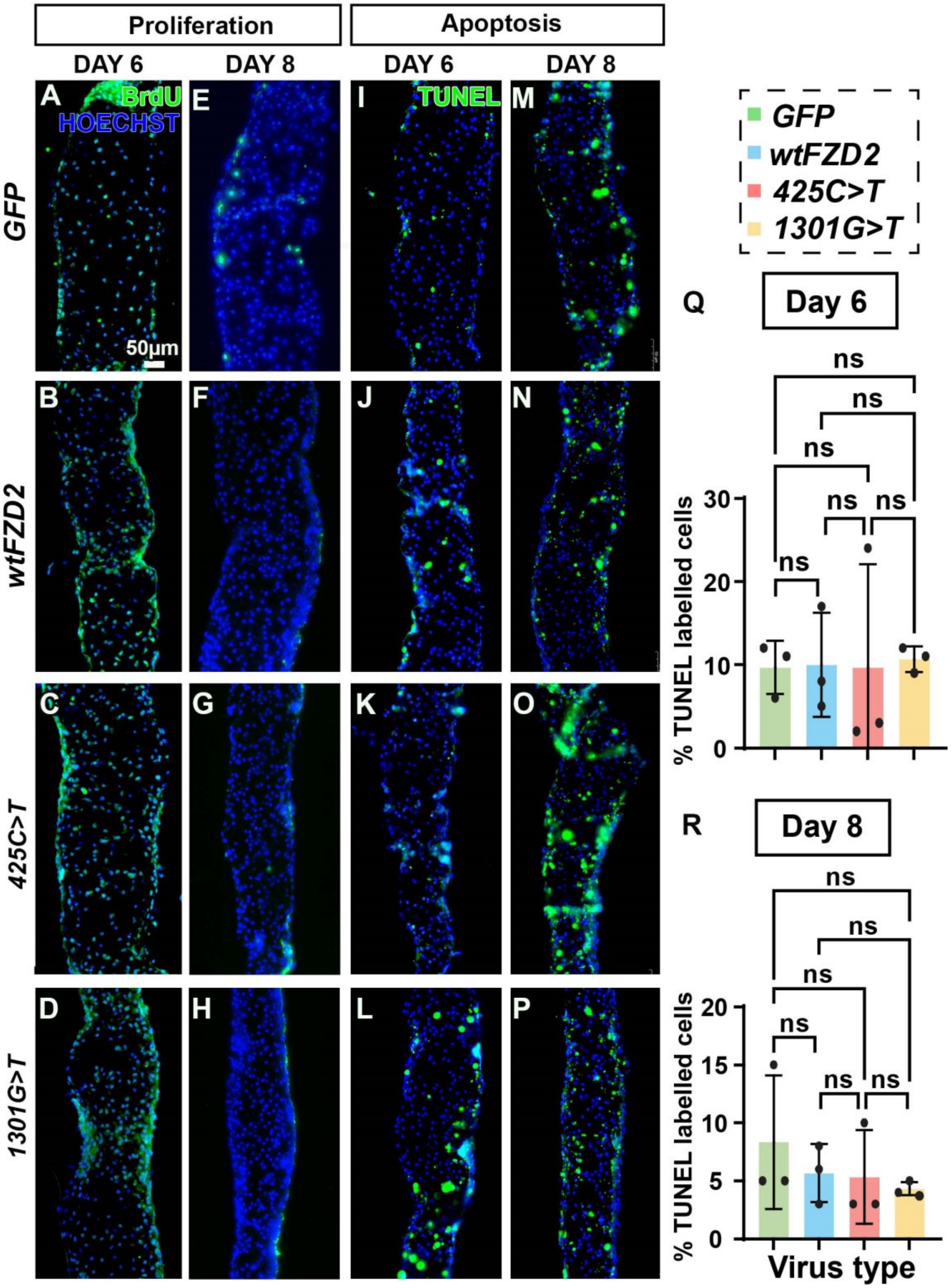

Proliferation
Apoptosis
DAY 6
DAY 8
DAY 6
DAY 8
A
BrdU
HOECHST
E
I
TUNEL
M
GFP
50μm
B
F
J
N
wtFZD2
C
G
K
O
425C>T
D
H
L
P
1301G>T
GFP
wtFZD2
425C>T
1301G>T
Q
Day 6
ns
% TUNEL labelled cells
30
20
10
0
R
Day 8
20
15
10
5
Virus type

**Figure S6: Distribution of proliferating and apoptotic chondrocytes in h*FZD2* infected day 6 and day 8 micromass cultures**

Serial sections from GAG-positive day 6 and day 8 micromass cultures were analyzed with anti-anti-BrdU (DSHB, G3G4) antibody. (A-D) On day 6, proliferating chondrocytes were found at the border of the cultures as well as in the cartilage forming area in all cultures irrespective of virus. (E-H) On day 8, the proliferating chondrocytes are located only at the border of the culture. TUNEL assay was performed on day 6 and day 8 cultures to detect the apoptotic cells in cultures. (I-L) On day 6 and (M-P) day 8 micromass cultures TUNEL positive cells were distributed throughout the culture in all viruses. (Q, R) Quantification of proportion of apoptotic cells showed no significant difference on day 6 and day 8 between GFP or hFZD2 viruses. Statistical comparisons between virus types were made using one-way ANOVA followed by Tukey's post hoc testing in GraphPad Prism 10.1.0. The error bars represent one standard deviation. Key: ns-not significant. Scale bar A-P = 50μm

**Figure S7**

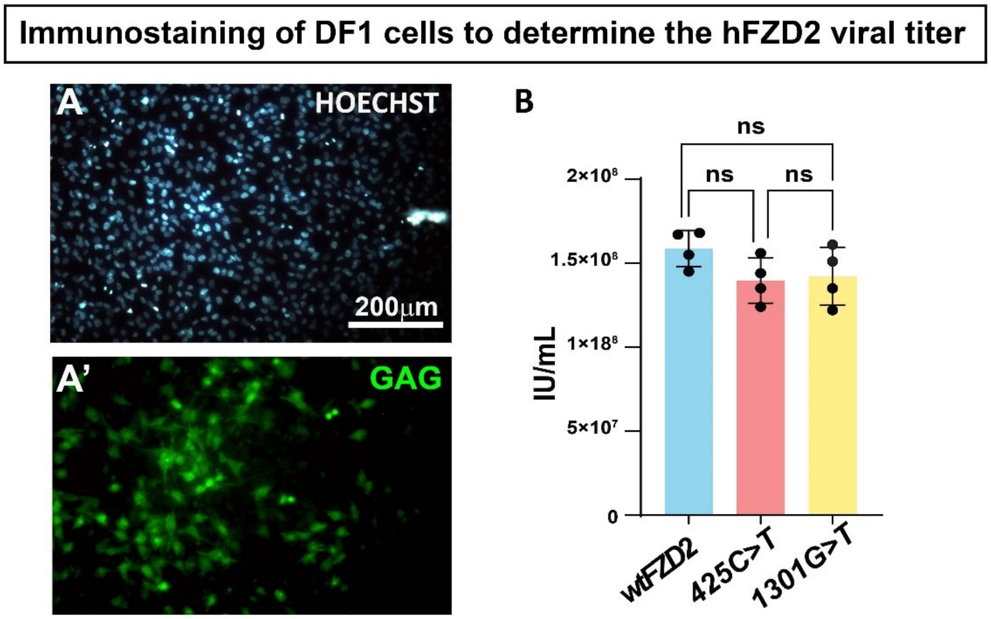


**Figure S7: Immunostaining on DF1 cells infected with high titer h*FZD2* virus and quantification of viral titer**

(A, A') Immunocytochemistry performed on DF1 chicken fibroblasts infected with 2μl of high titre viral stock were fixed at 36h post-infection. The cells were stained with anti-GAG antibody (virus) and counterstained with Hoechst. (B) Quantification of viral titer showed that all h*FZD2* viruses had higher than the recommended titer of 1 x $10^7$IU/mL for performing overexpression studies. The error bars represent one standard deviation. Scale bar = 200μm. Key: IU – Infectious units, ns – not significant

### Supplementary Tables

**Table S1: List of WNT pathway genes associated with Robinow Syndrome (White et al., 2015, 2016, 2018, Zhang et al. 2022)**

| Gene name | Inheritance pattern | Function | Variant type | OMIM |
|---|---|---|---|---|
| ***WNT5A*** | Autosomal dominant | Ligand | Missense | #180700 |
| ***DVL1*** | Autosomal dominant | Adaptor protein | -1 frameshift | #616331 |
| ***DVL2*** | Autosomal dominant | Adaptor protein | +1 frameshift | No entry |
| ***DVL3*** | Autosomal dominant | Adaptor protein | -1 frameshift | #616894 |
| ***FZD2*** | Autosomal dominant | Receptor | Missense, nonsense | No entry |
| ***ROR2*** | Autosomal recessive | Receptor | Missense, nonsense | #268310 |
| ***NXN*** | Autosomal recessive | Stabilizer of *DVL* | Deletion, missense | No entry |

**Table S2: ADRS associated missense and truncating *FZD2* variants reported in Zhang et al., 2022**

| | Total # of individuals | Phenotypes |
|---|---|---|
| **Total ADRS-FZD2 individuals from (Zhang et al., 2022)** | 17 | |
| **Missense mutations** | 7 | Broad forehead 7/7<br>Midface hypoplasia 7/7<br>Hypertelorism 7/7 |
| **Truncating mutations** | 9 | Broad forehead 6/7<br>Midface hypoplasia 5/7<br>Hypertelorism 2/7 |

**Table S3: Analysis of upper beak skeleton at stage 38 (E12.5) skulls stained in wholemount**

| RCAS Virus type injected at stage 15 (E2.5) | Stage 38 (E12.5) | External beak defects (N) | Internal bone phenotypes (N) | Missing bones | | | Reduced bone stain | | |
|---|---|---|---|---|---|---|---|---|---|
| | | | | pmx | n | prf | pmx | n | prf |
| ***GFP*** | 21 | 0 | 0 | 0 | 0 | 0 | 0 | 0 | 0 |
| **wild-type h*FZD2*** | 15 | 1 | 12 | 1 | 6 | 12 | 9 | 4 | - |
| ***425C>T*** | 13 | 1 | 9 | 1 | 4 | 9 | 1 | 3 | - |
| ***1301G>T*** | 19 | 6 | 16 | 2 | 7 | 16 | 5 | 4 | - |

**1** – Total embryos analysed for skeletal phenotypes caused by the FZD2 viruses
**2**- Total number of embryos with external phenotypes include short or deviated upper beak observed externally
**3** – The total number of samples with a phenotype were analyzed in detail and classified as embryos with missing bones or faint or absent alizarin red staining in bones derived from the frontonasal mass
**Key:** n; nasal bone, N; total sample size, pmx; premaxillary bone, prf; prefrontal bone, - missing bone, therefore ossification changes are not quantified

**Table S4: Survival of in vivo injected embryos**

| Virus type | Total injected embryos | Total embryos survived to day 10 | % survival |
|---|---|---|---|
| ***GFP*** | 42 | 31 | 73% |
| **wild-type h*FZD2*** | 36 | 23 | 64% |
| ***425C>T*** | 41 | 25 | 61% |
| ***1301G>T*** | 42 | 26 | 62% |

**Table S5: Percentage of BrdU and TUNEL positive cells in the frontonasal mass at stage 28 (3 days post-injection)**

| Virus type | Total injected | Total survived to day 3 (stg. 28) | Total embryos expressing GAG in frontonasal mass | % BrdU positive cells in the prenasal cartilage area (stg. 28) | % TUNEL positive cells in the prenasal cartilage area (stg. 28) |
|---|---|---|---|---|---|
| ***GFP*** | 8 | 8 | 7 | 6% | 2% |
| **wild-type h*FZD2*** | 9 | 9 | 6 | 7% | 3% |
| ***425C>T*** | 12 | 12 | 6 | 5% | 2% |
| ***1301G>T*** | 11 | 9 | 8 | 9% | 2% |

**Table S6. Restriction free Primer sequence for cloning *FZD2* variants into human wt *FZD2***

| ***FZD2* constructs** | **Primer sequence (5'→ 3')** |
|---|---|
| ***425 C→T*** | Fw: GAGCACTTCCTGCGCCACG<br>Rev: CGCGGCCCGATGGTTCC |
| ***1301 G→T*** | Fw: CCTCCTGGCCGTCTTCGTGTCGCTC<br>Rev: GCACTACACGCCGCGCATGTCG |

**Table S7: Micromass cultures used for histology and immunostaining analysis**

| **Virus type** | **Total cultures collected for histology/immunostaining** | | |
|---|---|---|---|
| | **Day 4** | **Day 6** | **Day 8** |
| ***GFP*** | 3 | 7 | 6 |
| **wild-type h*FZD2*** | 3 | 7 | 4 |
| ***425C>T*** | 3 | 9 | 4 |
| ***1301G>T*** | 3 | 7 | 6 |

**Table S8. Alkaline Phosphatase stain for staining mineralized fibroblasts in micromass**

| **Ingredient name** | **Distributor name, Product #** | |
|---|---|---|
| **Naphthol AS-MX phosphate** | Sigma Aldrich; N48751 | 1mg/ml dissolved in dimethyl sulfoxide |
| **Fast red violet LB salt** | Sigma-Aldrich; F3381 | 6mg/ml |
| **1M Tris (pH 8.3)** | Sigma; 77-86-1 | 10% |

**Table S9: Antibodies and immunofluorescence reagents**

| Antigen Retrieval (steam 15 minutes, 95°C) | Permeabilization/ Pre-treatment | Blocking serum | Primary antibody, source, dilution (Overnight, 4°C) | Secondary Antibody and counterstain |
|---|---|---|---|---|
| 10mM Sodium Citrate | 0.1% Triton X-100 in 1XPBS for 15 min | | **Ctnnb1/PY489-ß-catenin** Developmental Studies Hybridoma bank (DSHB), 5µg/ml, Mouse monoclonal | ThermoFisher, Cy5 goat anti-mouse, #A10524, ThermoFisher, Cy5 goat anti-rabbit, #A10523, |
| 10 mM sodium citrate | 0.5% hyaluronidase in Hank's Balanced Salt Solution, 45min (following antigen retrieval) | | **Collagen type II** DSHB, #II-II6B3, 5µg/ml, Mouse monoclonal | ThermoFisher, Alexa Fluor™ 488 goat anti-mouse, A11029, |
| 10mM Sodium Citrate | 0.5% hyaluronidase in Hank's Balanced Salt Solution, 45min (following antigen retrieval) | | **Collagen type II** Proteintech®, 28459-1-AP, 1:50, Rabbit polyclonal | ThermoFisher, Alexa Fluor™ 488 goat anti-rabbit A11034, 1:200 at RT in dark, 90mins |
| 1X Diva Decloaker (BioCare Medical) | 0.1% Triton X-100 in 1XPBS for 15 min | 10% Goat serum (Sigma G9023), 0.1% tween-20 in 1X PBS, (90 minutes, RT) | **Gag-pro AMV-3C2** DSHB, 1:4, Mouse monoclonal | |
| 10mM Sodium Citrate/ 1X Diva Decloaker (BioCare Medical) | 0.1% Triton X-100 in 1XPBS for 15 min | | **SOX9** Sigma-Aldrich, #HPA001758 1:200, Rabbit Polyclonal | **Hoechst** 10µg/ml (Sigma 33568) in 1 X PBS (RT in dark for 30 min) |
| 10mM Sodium Citrate/ 1X Diva Decloaker (BioCare Medical) | 0.1% Triton X-100 in 1XPBS for 15 min | | **TWIST1** Twist2C1a Abcam #ab50887 1:50, Mouse monoclonal | *Coverslip with ProLong™ Gold Antifade mountant (Invitrogen) |
| 0.3% Triton X-100 in 1XPBS for 15 min | 0.1% Triton X-100 in 1XPBS for 15 min | | **Flag (DYKDDDDK)** Thermo Fisher PA1-984B, 1µg/ml, Rabbit Polyclonal | |
| 1X Diva Decloaker (BioCare Medical) | 0.1% Triton X-100 in 1XPBS for 15 min | | **BrdU** DSHB, G3G4, 5µg/ml, Mouse monoclonal | |

**Table S10: qRT-PCR primers**

| Gene | Accession # | Forward Primer | Reverse Primer |
|---|---|---|---|
| ***hFZD2*** | NM001466 | CTTCCTGTGCTCCATGTACG | GCCACTGAAAACCGAACTTG |
| ***gFZD2*** | NM204222.2 | CTCCCATTGAGAGGACAGGT | TGTTTCGTTGTCCAGTCCAT |
| ***gDVL1*** | XM 015297120.1 | CTCCCATTGAGAGGACAGGT | TGTTTCGTTGTCCAGTCCAT |
| ***gWNT5A*** | NM003392 | CAATGGCTTCTCAGTACCTCG | ACATCTGCACAGGGTTCATG |
| ***gCTNNB1*** | NM 205081 | CTTGGACTTGACATTGGTGC | CAGAGTGGAAAGAACGGTAGC |
| ***gROR2*** | NM 001080716.1 | AGTGCTGGAATGAATTTCCC | GGGAACTTGTTTGTGTGGTG |
| ***gLRP5*** | NM001012897 | ACCAAAGCCAGAACCCAG | CAGCACCATCCCTATTGACTC |
| ***gATF2*** | NM 204904 | GCCAGCGTTTTACCAATGAG | AGTTGGTGTTGGTGTCTGATC |
| ***gLEF1*** | NM_001398055.1 | CATCAAGTCCTCGCTGGTC | GCCCTTGTCATGGTAGGAATC |
| ***gCOL2A1*** | NM 204426.1 | GGACCAGCAAGACGAAAGAC | CGTAGCTGAAGTGGAAACCG |
| ***gSOX9*** | NM_204281.2 | CTGGGCAAGCTGTGGAG | GGTTGGTACTTGTAGTCGGG |
| ***gTWIST1*** | NM_204739 | GACTCCAAGATGGCAAGCTG | CTCCATTCTCCACACCGAGA |
| ***gTWIST2*** | NM_204679 | GAGTTATGCCTTCTCAGTCTGG | ACGTCCCAATTCCACTTCAG |
| ***gBMP2*** | NM204358 | GCTGTTTTGAGGTGGATTGC | AGGCACTGTTCTCTTTGTCC |
| ***gBMP7*** | XM 417496 | GTCAAACATCGCAGAGAACAG | TCCTTCACAGTAATACGCAGC |
| ***gNOGGIN*** | NM204123 | ACTTTATGGCTATGTCCCTGC | AACTCCAGCCCCTTGATTTC |
| ***gBMPER*** | NM 001007080 | AGCTGTCCTCATGGTAAAATCC | AACGTACTGACATGTTCCCTG |
| ***gSP7*** | XM015300329.3 | GTTCGTCTGCAATTGGCTCT | AATTTCTTCTCGCGGGTGTG |
| ***gRUNX2*** | NM204128 | ACCTAGTTTGTTCCCTGAACG | GTAATCTGACTCTGTCCTTGTG G |
| ***gMSX1*** | NM_204559 | TTCGGTCAAATCGGAGAACTC | TTCGTCTTGTGCTTCCTCAG |

White, J. J., Mazzeu, J. F., Coban-Akdemir, Z., Bayram, Y., Bahrambeigi, V., Hoischen, A., . . . Mendelian, B.-H. C., 2018. WNT Signaling Perturbations Underlie the Genetic Heterogeneity of Robinow Syndrome. American Journal of Human Genetics. 102, 27-43.
Zhang, C., Jolly, A., Shayota, B. J., Mazzeu, J. F., Du, H., Dawood, M., . . . Carvalho, C. M. B., 2022. Novel pathogenic variants and quantitative phenotypic analyses of Robinow syndrome: WNT signaling perturbation and phenotypic variability. HGG Adv. 3, 100074.
